# Adding layers of information to scRNA-seq data using pre-trained language models

**DOI:** 10.1101/2025.08.23.671699

**Authors:** Sonia Maria Krißmer, Jonatan Menger, Johan Rollin, Tanja Vogel, Harald Binder, Maren Hackenberg

## Abstract

Pre-trained language models promise to enrich single-cell analyses with contextual information from large biomedical text corpora, but it remains unclear how to optimally align this knowledge with quantitative scRNA-seq data. To address this, we construct text-based training datasets from both scRNA-seq data and biomedical literature targeted to the experimental setting at hand. We then fine-tune lightweight encoder-only biomedical language models to learn a shared, literature-enriched representation. Controlled evaluations across immune and developmental datasets show that this representation preserves cell identity while adding robust and interpretable contextual layers of functional, disease-associated, and developmental information to single-cell analysis workflows.

## 1 Introduction

In single-cell data analysis, an increasing number of foundation models pre-trained on a large amount of data has recently become available [1]. This includes models trained directly on quantitative single-cell profiles (e.g.,[2, 3]), and biomedical language language models trained on large text corpora, (e.g.,[4, 5, 6]). While expression-based models learn structure from molecular profiles [2, 7, 3, 8], language models that leverage biomedical literature can encode complementary contextual information about genes, cell types, diseases, and developmental processes.

Most current applications investigate both types of foundation models as primary end-to-end alternatives to classical quantitative single-cell analysis workflows, for tasks such as cell type annotation, batch integration, or perturbation prediction [9, 10, 11, 12, 13]. A complementary approach is to use them to add contextual layers to the original data and augment standard analyses. In such a setting, language models trained on scientific literature can be used to embed cells together with qualitative biological descriptors, such as functional programs, disease associations, or developmental stages. Yet, it remains unclear how to best leverage such textual knowledge to enrich analyses of a specific dataset of interest [13].

A natural approach for such integration is to convert single-cell profiles to “cell sentences” [10, 14], which represent each cell as a ranked list of its most highly expressed genes. This text-based representation makes it possible to combine expression-derived information with dataset-specific metadata and related contextual information retrieved from scientific literature in a common format. However, this also raises a methodological challenge. Text-and expression-derived information need to be combined in a biologically meaningful and interpretable way, so that single-cell profiles, metadata, and literature-derived concepts can be compared within the same representation. Further, because cell type labels, experimental metadata, and literature search terms can each shape the learned embedding, their individual contributions need to be evaluated explicitly to interpret downstream results. Here, we explore a contrastive alignment strategy to learn a joint embedding space in which expression profiles and literature-encoded knowledge are directly comparable, and use systematic ablation experiments to assess the contribution of each information source. Specifically, we train small encoder-only language models [15, 16], which allow for flexible, task-specific fine-tuning adapted to a given dataset, support straightforward integration of any text-based data source, and can generate semantically meaningful representations compatible with established downstream analyses.

Current research on pre-trained language models for single-cell data focuses mainly on large language models (LLMs) that can be employed for diverse primary analysis tasks such as cell type annotation, batch integration, and perturbation prediction. For example, C2S-Scale [10] trains generative language models such as Pythia [17] on cell sentences to support both standard single-cell tasks and dataset interaction via natural language queries. Other approaches use general-purpose LLMs to create cell embeddings based on gene identities or expression-weighted gene vectors [11, 7], or for cluster-based annotation [9]. These approaches show the flexibility of language-based representations for single-cell analysis. At the same time, it remains unclear when such large generalist models are necessary, and when smaller, dataset-adapted models are sufficient for targeted contextualization tasks [13].

A related set of approaches combine quantitative and text-based data representations and use language models for text-based interaction with the quantitative data. For example, CellWhisperer [18] aligns expression-based cell embeddings with textual descriptions and provides a chat-based interface to query cells in natural language. Similarly, models that combine transcriptomic foundation models such as Geneformer [3] or scGPT [2] with biomedical text encoders like BioBERT [19] integrate raw expression data with textual knowledge in a shared space [4, 18, 20]. However, scientific literature is typically incorporated only indirectly through general pre-training rather than as a targeted source of contextual knowledge specific to the dataset of interest [13]. In addition, when labels, metadata, and text-derived information are combined, their individual contributions to downstream results remains difficult to assess.

Therefore, we propose a targeted approach for incorporating biomedical literature into scRNA-seq analysis, focusing on smaller, dataset-adapted encoder-only language models for adding contextual layers of information to the original single-cell data. We first express all information sources in textual form with a consistent structure. We represent single-cell profiles as cell sentences, add metadata using sentence templates, and retrieve biomedical literature from PubMed [21] and additional biomedical databases using queries targeted to the experimental metadata. Each cell sentence is paired with a similar and a dissimilar sentence to form a triplet [22, 23], and corresponding triplets are constructed from PubMed titles and abstracts. This triplet structure enables contrastive fine-tuning of small sentence embedding models [15, 16] to align cell-derived and literature-derived texts in a shared semantic embedding space [24, 25].

We show that the resulting representation embeds text-based single-cell profiles, annotations, and literature-derived concepts within a shared coordinate system that can be queried across different biological layers. We use matched ablations and validation on held-out donors, mice, and developmental time points to clarify how expression-derived, metadata-derived, and literature-derived information shape the learned embeddings. Across immune and developmental datasets, the embeddings preserve cell identity and enable functional, disease-associated, and developmental analyses. Together, these results show that lightweight biomedical language models can be used as complementary tools for adding controlled and interpretable layers of contextual information to single-cell analysis workflows.

## 2 Results

### 2.1 A unified workflow to embed single-cell profiles in the context of biomedical knowledge

To integrate single-cell gene expression profiles and complementary biomedical knowledge, we developed a text-based approach that embeds cell-derived and literature-derived information into a shared semantic space (Fig. 1). In this representation, individual cells and biomedical concepts expressed in textual form (such as cell type labels, metadata descriptions, or literature-derived concepts) can be compared directly. This enables to add contextual layers such as function, disease association, or developmental state to the data.

**Figure 1:**
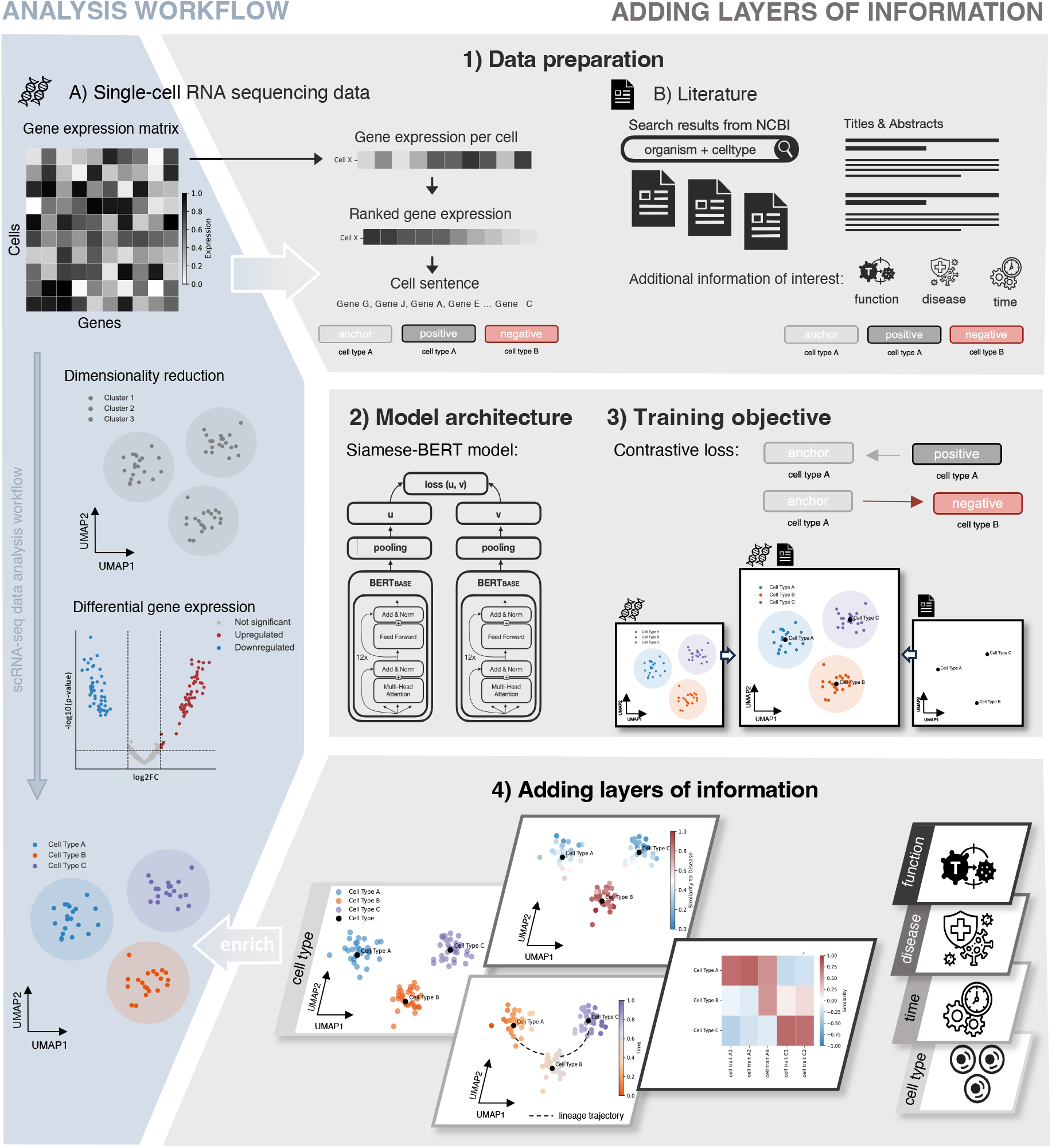
Overview of the approach for adding contextual layers of information to scRNA-seq datasets. Left: A conventional single-cell RNA-seq workflow processes a gene expression matrix through a quantitative pipeline comprising dimensionality reduction and differential expression analysis for annotation of cell type clusters. **1)** In our approach, we translate single-cell profiles into cell sentences by ranking highly expressed genes and appending metadata for generating the cell-derived dataset (A). In parallel, we build a literature-derived dataset by retrieving PubMed titles and abstracts related to the known or suspected cell types, capturing complementary information such as cell type functionality (*function*), disease relationships (*disease*) or lineage trajectories (*time*). For both datasets, we construct label-aware triplets. **2)** On the triplets, we train models with a Siamese-BERT [16] architecture, using **3)** contrastive fine-tuning to align the embeddings of single cells, cell type labels and literature-derived information. **4)** The resulting joint embedding space enables context-aware analysis of scRNA-seq data with additional layers of biological information captured in the biomedical literature.

Our approach starts from scRNA-seq datasets processed by a conventional analysis workflow (Fig. 1 left) with annotated cell types, e.g., based on cluster-specific differential gene expression analysis [26, 27]. We use such annotations as dataset-specific prior information for constructing training examples. Starting from the gene expression matrix, we transformed each cell’s gene expression profile into a cell sentence by ranking highly-variable genes according to their expression and concatenating the top-ranked gene symbols. A supplementary sensitivity analysis on a smaller PBMC dataset showed that cell sentences of 50 genes still retain sufficient structure while keeping sentence length computationally manageable, with limited variability across the tested random seeds (Additional file 1, Fig. S1), and we thus used cell sentences of length 50 as a pragmatic choice for all analyses. Depending on the downstream task, we added metadata such as cell type, donor, or developmental time point using sentence templates (Fig. 1, Step 1 (top left)).

In parallel, we built a literature-derived training dataset by retrieving titles and abstracts of research articles stored in the National Center for Biotechnology Information (NCBI) PubMed database [21] that are relevant to the organism, cell types, or developmental stages represented in the scRNAseq data (Fig. 1, Step 1 (top right)) as well as cell ontology cell type descriptions [28] and known marker genes [29] were available. This allows for capturing complementary biological knowledge not directly annotated in the gene expression profiles, such as cell type functionality, disease relationships, or developmental trajectories. We thus obtained a gene expression-derived and a literature-derived dataset in textual format, facilitating direct embedding by a language model into a shared space.

To align embeddings of both datasets based on their semantic relationship, we used a label-aware triplet strategy for fine-tuning a language model. Within the expression-derived and the literature-derived datasets, we constructed triplets consisting of an anchor, a positive sample sharing the same cell type label, and a hard negative sample with a different label but high embedding similarity (Fig. 1, Step 1, bottom; Methods Section 5.6). On these triplets, we fine-tuned a small encoder-only language model with a Siamese-BERT architecture [16] (Methods Section 5.5), designed to efficiently compute similarity of multiple inputs (Fig. 1, Step 2). Training alternates epoch-wise between the cell-derived and literature-derived training data (Methods Sections 5.8) and employs a multiple negatives ranking loss function, designed to capture semantic relationships within triplets by pulling positive samples towards the anchor while pushing negative samples further apart. The resulting joint embedding space is therefore optimized to place biologically related cell sentences, annotations, and literature-derived texts close to each other while separating unrelated examples.(Fig. 1, Step 3; Methods Section 5.9). After fine-tuning, we visualized the learned embedding using UMAP [30]. Taken together, the approach allows for enriching representations of single-cell profiles by adding layers of complementary contextual information, such as cell type, function, disease, or time (Fig. 1, Step 4).

We evaluated the workflow using cell sentences from held-out donors or mice without metadata labels throughout, and held-out time points in the developmental example (Methods Sections 5.1, 5.2; Additional file 2, Fig. R1). Because labels, metadata, and literature-derived search terms can each shape this representation, we trained matched ablation models to assess the respective contributions of expression-derived, metadata-derived, and literature-derived information (Methods Section 5.7). After assessing whether embeddings preserve known cell identities, we illustrate how the learned embeddings allow for annotation based on functional descriptions not used during training, characterization of disease-associated functional shifts, and developmental trajectory analysis.

### 2.2 Text-based fine-tuning preserves cell identity in controlled embedding representations

We first assessed whether contrastive fine-tuning provides embeddings that preserve cell identity while aligning cell-derived sentences with textual cell type information as a prerequisite for using the representation as a contextual layer on top of standard scRNA-seq analyses. The embedding should retain known cell type structure, allow for label-based annotation, and be robust to querying with paraphrased descriptions instead of the label used during training.

We used a T cell subset of the Human Immune Health atlas from the Allen Institute (HIAI dataset, Methods Section 5.1) [34]. As an initial qualitative assessment, we compared UMAP representations of the HIAI T-cell dataset obtained from the base language model before fine-tuning, a model trained only on literature-derived text, and the full model trained on both cell-derived and literature-derived datasets (Fig. 2 **A**). The base model showed poor separation of cell subtypes, whereas joint finetuning resulted in an embedding with distinct cell clusters by subtype, with their respective cell type label embedded within those clusters. We also observed meta-clusters of CD4+ and CD8+ T cells and a tendency of cells with similar functionality to also cluster together in the embedding, such as memory CD8+ T cells and γδ T cells, both characterized through strong cytotoxic capabilities [35, 36]. This is supported by assessment of marker gene expression in the model embeddings (see Additional file 1, Fig. S2), confirming that the model embeddings capture biologically meaningful marker gene expression patterns while preserving within-cell-type heterogeneity.

**Figure 2:**
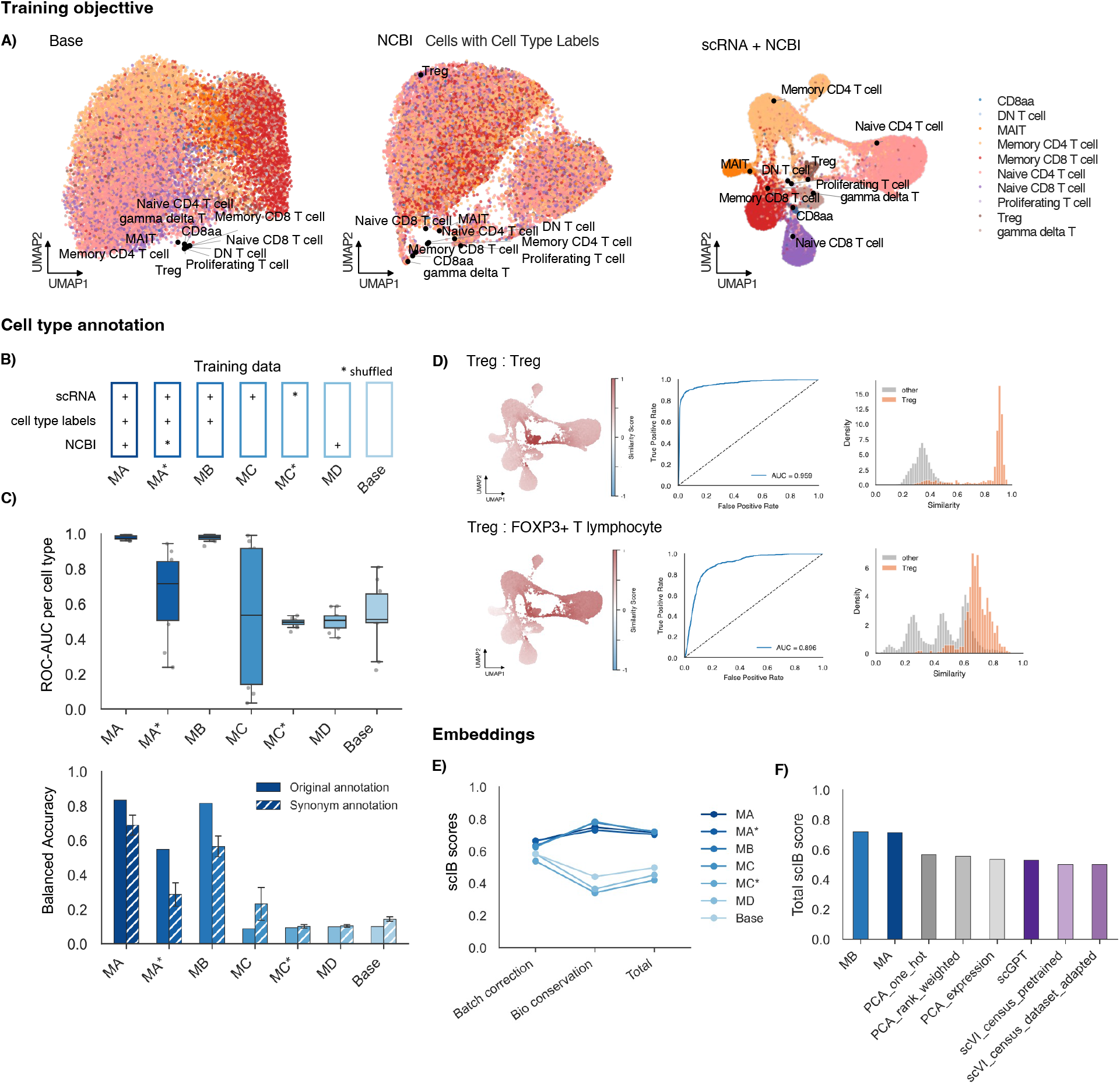
Joint fine-tuning on cell-derived and literature-derived datasets preserves cell identity in joint text-based embeddings of the HIAI T-cell dataset. **A)** UMAP representation based on embeddings from the pre-trained base model (left), a model trained only on literature-derived data (middle), and the full model trained on cell-derived and literature-derived datasets (right). Cells are colored by the reference cell type annotation from the HIAI dataset; embedded cell type labels are shown in black. **B)** Ablation design. Columns indicate model variants and rows indicate training information sources. Plus signs indicate included sources; stars indicate shuffled labels for the corresponding source. MA: full model including scRNA-seq-derived cell sentences, cell type labels, and NCBI-derived literature; MA*: full model with shuffled literature labels; MB: semantic cell sentences with cell type labels; MC: cell sentences only, using the correct cell type for building triplets; MC*: cell sentences only, using shuffled cell types for building triplets; MD: literature-derived text only; Base: pre-trained model without fine-tuning. **c)** cell type label alignment and annotation performance across ablation models. Top: Boxplots show one-vs-rest ROC-AUC values across cell types based on cosine similarity between cell embeddings and cell type label embeddings. Bottom: Bar plots show overall balanced accuracy after assigning each held-out cell to the cell type label with highest cosine similarity, using both the original annotation (solid bars) and synonym annotations (striped bars). **D)** Distributions of cosine similarity scores distributions in the learned embedding (left), ROC curves (middle), and score histograms (right) for regulatory T cells queried with the original annotation label (‘Treg’) and a paraphrased description (‘FOXP3+ T lymphocyte’). **E)** Embedding quality of ablation models evaluated with single-cell integration metrics [31] for batch correction, biological conservation, and total score. **F)** Total integration benchmark scores for MA and MB models and reference approaches (PCA variants, scGPT, variants of scVI [32] pre-trained on a CELLXGENE dataset collection [33].

To determine how each training sources contributes to this alignment, we then trained matched ablation models varying the inclusion of cell sentences, cell type metadata, literature-derived text, and including shuffled controls (Fig. 2 **B**: Methods Section 5.7). We quantify cell identity preservation using label-alignment and annotation metrics (Fig. 2 **C**; Methods Section 5.10): We used label-alignment ROC-AUC to evaluate for each cell type how well cosine similarities of cell profile embeddings to the cell type label embedding separate cells from that type from all others. In addition, we used balanced accuracy to evaluate multiclass cell type annotation after assigning each cell to the label with the highest similarity, using both the original labels and synonym annotations. Comparing all ablation models shows that cell type annotation and cell type label alignment is largely driven by correctly structured label information in the cell-derived training data, and shuffling literature information breaks the alignment even if correct labels are present. The similar performance of the full model and the model trained without literature for many cell types indicates that literature-derived text does not uniformly improve basic cell type annotation, but rather contributes in a class-and query-dependent manner (Additional File 1, Fig. S3 **A**-**D**). Finally, Fig. 2 **C** (bottom) shows that literature-derived information is particularly important for synonym-based annotation. To further contextualize these results, we report cell type abundances in the scRNA-seq-derived data and retrieval counts for the literature-derived data (Additional file 2, Tables R1-R4).

As an external reference, we compared our results with CellTypist [37, 38] and SingleR [39], because both methods could be trained or configured on the same donor-level training data split and thus enable a matched comparison of dataset-adapted annotation performance (Additional file 1, Fig. S3 **A, B**; Methods Section 5.11). The full model achieved a balanced accuracy of 83.3 %, which is slightly below fine-tuned CellTypist and above SingleR. The approach thus provides competitive annotation performance on held-out donors, while not being optimized or primarily intended as a cell type annotation tool.

For the full model, we additionally visualized similarity scores between cell embeddings and the embeddings of exemplary candidate cell type labels and compared similarity score histograms and ROC curves (Fig. 2 **D**). For regulatory T cells, similarity to both the original label (‘Treg’) and a synonym (‘FOXP3+ T lymphocyte’) separated the corresponding cells from other T-cell subsets (Fig. 2 **D**), indicating that cell type alignment is not restricted to exact string matching.

We observed lower prediction accuracy within functionally similar cell types, such as γδ T cells and memory CD8 T cells, or regulatory T cells with other CD4+ T-cell subtypes [35], reflecting biologically plausible overlap between closely related functional states (see Additional file 1, Fig. S3 **C**, **D**).

Finally, we assessed the models’ embedding quality using standard single-cell integration bench-marking metrics for biological conservation and batch correction [31](Fig. 2 **E**, **F**; Additional File 1, Fig. S3 **E**, **F**; Methods Section 5.12). The ablation models show that while correctly structured labels in the cell-derived data primarily drive cell type annotation and literature-derived information improves synonym annotation, the embedding quality primarily depends on the inclusion of non-corrupted scRNA-seq-derived data (Fig. 2 **E**). Models with non-shuffled labels and labels+literature information achieve higher overall scores than baselines (Fig. 2 **F**).

Together, these analyses support that the training strategy preserves cell identity and that semantic label alignment depends on correctly structured metadata and literature, and provides a controlled basis for evaluating downstream contextual layers.

### 2.3 Literature-aligned embeddings capture functional programs at single-cell level

Next, we asked whether the text-based embeddings also capture functional information beyond cell type labels. On the HIAI T cell dataset, we curated descriptions of T-cell functionality (see Additional file 1, Figure S4 **A**) and evaluated whether these descriptions could be assigned to the corresponding cell types or cells in the learned embedding space.

For each expert-curated functionality description, we ranked all candidate T-cell types according to their association with the description on two levels (Methods Section 5.13.1). At the cell-type-label level, we computed cosine similarity between the embedded descriptions and the embedded cell type labels. At the cell level, we computed similarity scores for each individual cell and ranked cell types according to the cell-level scores. We then summarized performance using mean reciprocal rank (MRR) across cell types. The full model achieved the highest MRR, with best performance for cell-level assignment (Fig. 3 **A**). In contrast, models without correctly aligned literature performed worse, while the literature-only and base models retained some performance mainly at the cell-type-label level. This indicates that while general biomedical language representations encode relationships between cell-type names and functional descriptions, explicitly aligning literature-derived text with cell-derived sentences is necessary to transfer this information to single-cell embeddings. This stronger dependence on literature-derived text contrasts with basic cell type annotation, where correctly structured cell-derived label information already accounts for much of the performance.

**Figure 3:**
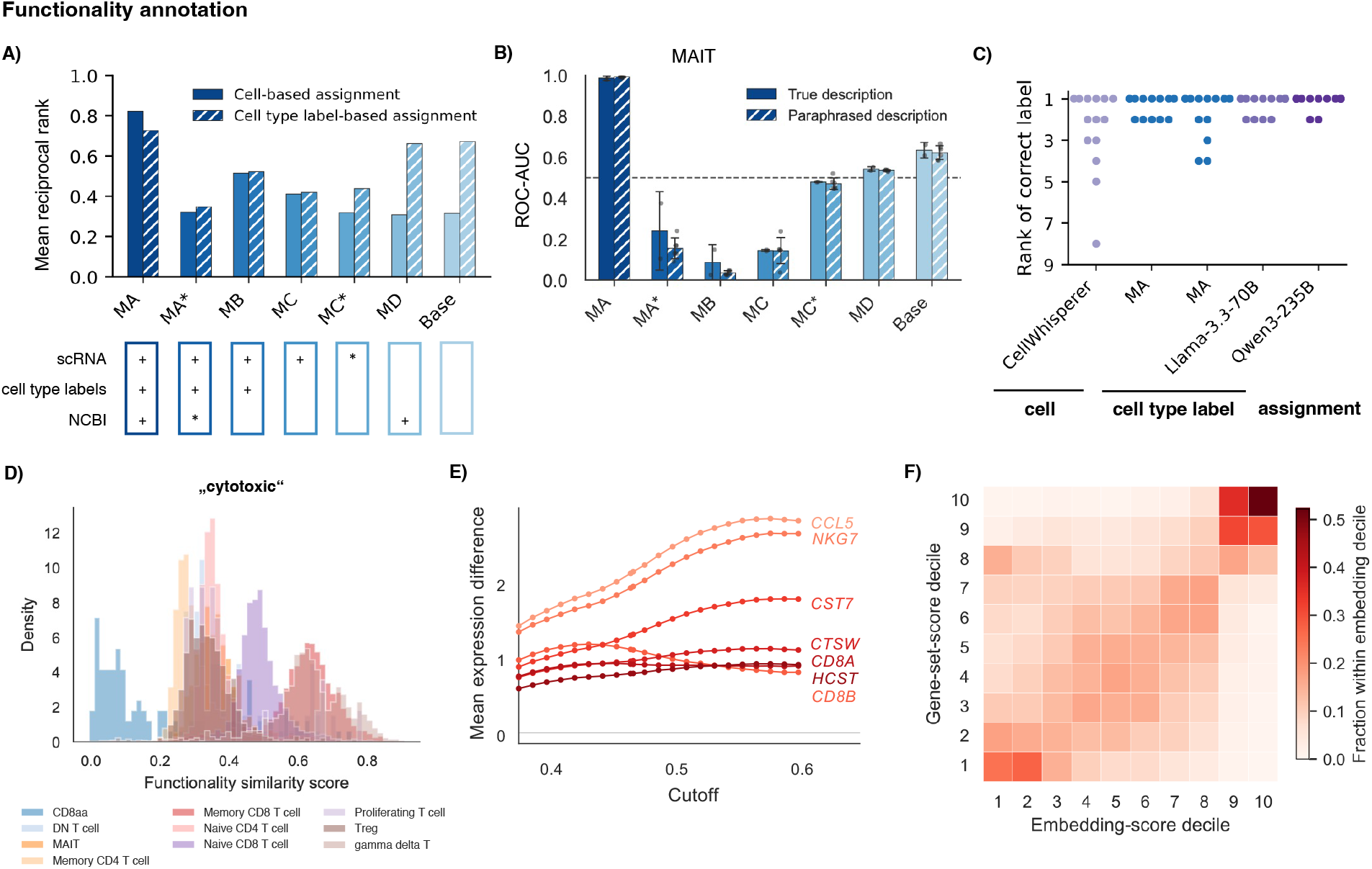
Literature-aligned embeddings capture functionality T-cell programs at the cell-type and single-cell level. **A)** Functional assignment across ablation models, summarized by mean reciprocal rank (MRR). For each expert-curated functionality description, candidate T-cell types were ranked by association with the description; higher MRR indicates that the expert-assigned cell type was ranked closer to the top. Solid bars show assignment based on single-cell embeddings, striped bars show assignment based on cell type label embeddings. **B)** Robustness to paraphrasing for MAIT-cell functionality. ROCAUC values are shown across ablation models for the original expert-curated MAIT-cell functionality description (solid bars) and three paraphrased descriptions (striped bars). **C)** Comparison of the full encoder-only model with existing generalist and specialist models on the single-cell and cell-type label-based level, using rank-based association of functionality descriptions with expert-assigned cell type labels or cells. **D)** Example application: histogram of cell-wise similarity scores to the functionality annotation ‘cytotoxic’, colored by reference cell type label, showing separation of cells with known cytotoxic identity. **E)** Sensitivity of differential expressed results of cells annotated as cytotoxic vs non-cytotoxic cells to the decision threshold, confirming robust identification of known cytotoxicity markers. **F)** Concordance between embedding-based cytotoxicity similarity scores and cytotoxicity scores from an established gene set after binning cells into score deciles.

Repeating the analysis for MAIT cells using the original expert-curated description and several paraphrased versions showed that the full model retained high ROC-AUC for both formulations, whereas ablation models showed weaker or less consistent performance (Fig. 3 **B**).

When investigating results for individual descriptions, we observed higher similarity between functionality descriptions that share biological context, for example between memory-associated T cell subtypes or CD4+ in contrast to CD8+ T cells (see Additional file 1, Fig. S4 **B**).

Next, we compared the full model with existing single-cell foundation models and general-purpose generative language models using the same cell-based and cell-type-based comparison strategies (described in Methods Section 5.13.2). It performs slightly better than CellWhisperer [18] on the single-cell level, and performs comparable to or slightly worse than generalist language models on the cell type level (Fig. 3 **C**).

We then used the embedding to identify overarching functional programs that span multiple cell types (Methods Section 5.13.3). For the query’cytotoxic’, similarity scores separated a subset of CD8+ as well as γδ T cells with high functional similarity from the remaining T cells (Fig. 3 **D**). Using different thresholds around a cutoff of 0.468 determined via Otsu’s method [40], we assessed the stability of downstream differential expression results (Methods Section 5.13.4). The top genes identified at the automatic Otsu cutoff established cytotoxicity markers [41] and their expression differences are robust across thresholds, indicating that the functional query selected a biologically coherent cell population (Fig. 3 **E**).

Finally, we compared the text-based functional similarity scores with standard gene-set scores for cytotoxicity (Methods Section 5.13.5). Decile-based concordance analysis showed that cells with high embedding-based cytotoxicity scores also tended to have high cytotoxicity gene-set scores (Fig. 3 **F**, Additional file 1, Fig. S5 **F**, **G**). We observed analogous patterns for an additional ‘regulatory’ query (Additional file 1, Fig. S5 **A**-**E**), showing that text-based query results were broadly consistent with conventional marker-based summaries, while allowing functional programs to be specified in natural language.

### 2.4 Embeddings recover disease-associated functional shifts

To assess whether our approach can effectively incorporate both literature-derived knowledge and metadata, we focused on cytomegalovirus (CMV)-associated cytotoxic shifts in T cells and included CMV-related literature in the literature-based triplet construction. To avoid direct use of the target disease label, CMV status was not included in the cell sentences, and all evaluations were performed on held-out donors.

Fig. 4 **A** shows CMV status across all cells in the UMAP representation of the original scRNA-seq dataset. We computed similarity scores between each held-out cell embedding and the interaction description “increased cytotoxic activity in cytomegalovirus positive patients”. Within each cell type, we compared the distributions of cell-wise similarity scores between CMV-positive and CMV-negative donors using a Mann-Whitney U test (see Methods Section 5.14, 5.15). As cells from the same donor are not statistically independent, cell-level tests are used here as descriptive comparisons and not interpreted as donor-level inferential evidence. In Fig. 4 **B**, cells from CMV-positive held-out donors showed higher similarity scores within memory CD4+ T, memory CD8+ T and γδ T cells than cells from CMV-negative donors. In particular, CD4+ T cells are typically non-cytotoxic in healthy individuals but can acquire cytotoxic properties upon CMV infection [35], reflecting a known functional shift.

**Figure 4:**
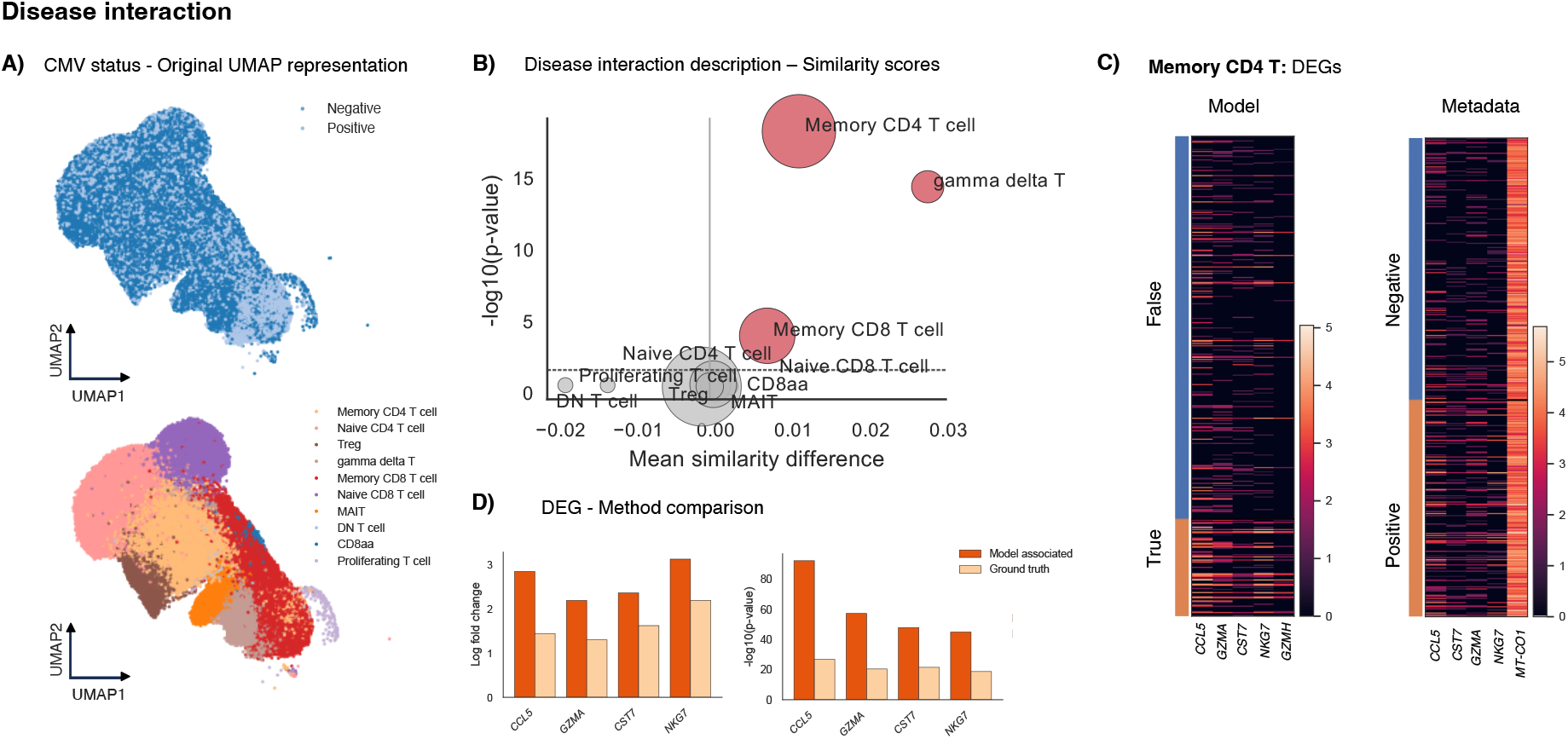
Integrating literature-derived knowledge and metadata enables detection of disease-associated functional alterations in cytomegalovirus (CMV)–positive individuals. **A)** UMAP representation of the original gene expression data annotated with CMV status metadata and cell type labels. **B)** Separation of CMV status based on cell-wise cosine similarity to a disease interaction description, where cells are classified as CMV-associated (“True”) when exceeding the similarity threshold (see Methods Section 5.14). Separation is evaluated for cells of separate cell types using the mean similarity difference and p-value from Mann-Whitney U test. Dot size indicates the number of cells per type. **C)** Differential gene expression (DEG) analysis of disease associated cells’Memory CD4 T’ cells separated either by similarity scores based on the model embeddings or by the metadata labels. **D)** Comparison of log fold changes (logFC) and corresponding log-transformed exploratory *p*-values between the two DEG analyses from the two methods of cell separation.

To further characterize the expression patterns associated with high CMV-related similarity scores, we performed differential gene expression analysis for memory CD4+ T cells using two alternative separations: the available CMV metadata and the model-derived similarity scores, where we used Youden’s J index [42] to determine an optimal threshold (Fig. 4 **C**, Methods Section 5.14). Four of the five most strongly up-regulated genes were shared between the two analyses. For all four shared genes, the model-derived grouping provided larger log-fold changes than the grouping based on donor-level CMV metadata (Fig. 4 **D**). Together, these results indicate that the model-derived scores recover a known CMV-associated cytotoxic expression pattern and enrich for cells showing this phenotype, despite CMV status not being included in the cell sentences. We obtained comparable results with a model trained without dedicated CMV-specific literature or CMV metadata, suggesting that relevant CMV-associated cytotoxicity information was also captured in broader cell-type-associated literature (Additional file 1, Fig. S6).

### 2.5 Embeddings capture developmental organization along lineage trajectories

We next evaluated whether our approach captures temporal metadata information and developmental dynamics. For this, we used an embryonic mouse brain dataset [43], containing a diverse range of cell types sampled at multiple time points of mouse embryonic development from day 7 to day 18. Temporal metadata (embryonic day) was included into the generated semantic cell sentences as described in Section 5.2, while evaluation was performed on cell sentences from held-out mice and not including metadata labels. To facilitate visualization, we divided the dataset into seven subsets (results for all subsets shown in Additional file 1, Fig. S7, S8). Here, we compared the alignment between the original authors’ cell type annotations (color code) and the model’s cell type predictions (cell label), along with the embryonic time shown in Additional file 1, Fig. S7, S8 **A)**. The fine-tuned model aligned the embeddings of cell type labels and their corresponding cells with a mean ROC-AUC of 0.905.

Beyond alignment of cell type labels and corresponding cell profiles, the embeddings reflected broad temporal organization in meta-clusters spanning multiple annotated cell types. Cells from early embryonic differentiation states clustered separately from cells of more mature states (Additional file 1, Fig. S7, S8 **B)**). This suggests that when incorporating temporal information, the model embedding captures both cell type identity and temporal patterns in held-out mice.

We then focused on a forebrain trajectory of cells from the ‘Dorsal forebrain’ to mature ‘Cortical and hippocampal glutamatergic’ neurons, a sub-lineage of one of the seven broader subsets (Fig. 5 **A**). We computed diffusion pseudotime on both the model embeddings and the original gene expression matrix (Methods Section 5.16). The ranked pseudotime values showed a strong concordance between the expression-based and model-based representations (Kendall’s *τ* = 0.631, − log_10_ *p* = 6595; Fig. 5 **B**). Both pseudotime estimates were broadly consistent with the chronological order of the embryonic days in UMAP representations of the model embeddings and the original gene expression matrix (Fig. 5 **C**). The model embeddings additionally separated a subset of *Crabp2*+, *Fezf1*+ and *Hes5*early forebrain neural progenitors previously annotated as ‘Forebrain’ [44, 45] (gene expression of the mentioned markers in Additional file 1, Fig. S9). This subset was consistent with early time points in the metadata annotation and early model-based pseudotime predictions, while the expression-based pseudotime did not separate these early neuronal progenitors. This suggests that the model embeddings retain developmental structure along this trajectory.

**Figure 5:**
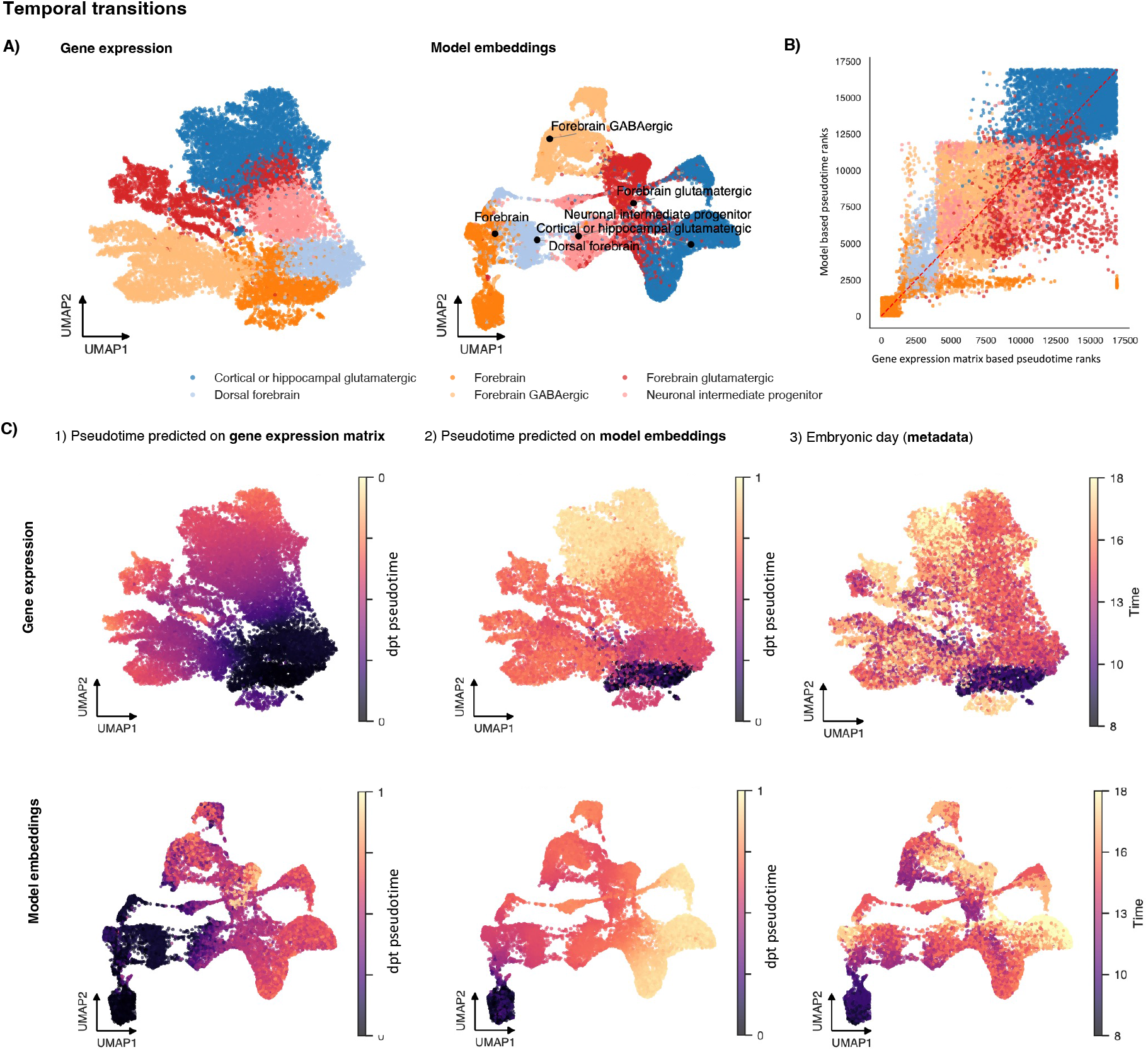
Model embeddings incorporating temporal metadata capture developmental ordering along a forebrain lineage. **A)** UMAP representations of cells from the trajectory from ‘Dorsal forebrain’ to ‘Cortical or hippocampal glutamatergic’ and ‘Forebrain GABAergic’ colored by the annotated reference cell type label, using the gene expression matrix (left) or the model embeddings (right). **B)** Concordance of diffusion pseudotime ranks computed from the gene expression matrix (x axis) and model embeddings (y axis) colored by the annotated reference cell type label. **C)** UMAP representations based on the gene expression matrix (top row) and the model embeddings (bottom row), colored by diffusion pseudotime values computed based on model embeddings (left), pseudotime values computed based on the gene expression data (middle), and embryonic day metadata (right).

As a supplementary qualitative validation, we withheld selected embryonic time points during training and projected cells from these time points into the learned embedding space. Cells from heldout time points mapped to developmentally plausible regions of the embedding, which is confirmed through neighborhood comparison with gene expression-based embeddings and supports coarse temporal generalization (Additional file 1, Fig. S10).

Together, these results indicate that temporal metadata and literature-informed training can capture continuous developmental transitions and broad hierarchical lineage relationships, and thus support developmental trajectory analyses.

## 3 Discussion

We have investigated how pre-trained encoder-only biomedical language models can be adapted to enrich classical scRNA-seq workflows with contextual information from biomedical literature, more generally exploring language models as a complementary analysis tool. Our approach converts cell profiles, metadata, and literature-derived information into a shared text-based representation and aligns them using contrastive fine-tuning. We have applied our approach to single-cell RNA-seq datasets from human immune and developing mouse brain data to evaluate its ability to integrate literature-derived context across biological systems. Our results have shown that the resulting embeddings provides a semantically meaningful, unified representation of information from these data sources and can connect gene expression profiles of individual cells with textual descriptions of their biological roles and function. They preserve cell identity while adding interpretable layers of functional, disease-associated, and developmental information. Matched ablation studies, held-out validation, and controls using paraphrased textual descriptions confirm generalizability and robustness of the embeddings and allow to assess what each information source uniquely contributes.

In the HIAI T cell dataset, the embeddings preserved cell type structure and achieved competitive annotation performance on held-out donors compared with dataset-adapted baselines, although the model is not primarily designed as an annotation tool. Ablation analyses showed that non-corrupted scRNA-seq-derived input is central for embedding quality, whereas correctly structured labels and literature-derived text are important for semantic label alignment and functional assignment. Cell type functionality descriptions and functional queries such as cytotoxicity and regulatory activity were robust to paraphrasing and annotation threshold choice and coherent with conventional gene-set scores. When investigating disease associations, the embeddings reflected phenotypic alterations in CMV-positive individuals, recovering known cytotoxic shifts in CD4+ T cells in held-out donors without including of CMV status in the training data. Comparable results without dedicated CMV-specific literature suggest that relevant disease-associated context can also be present in broader cell-type-associated literature. In a developing mouse brain dataset, the inclusion of temporal metadata enabled the model to reconstruct developmental progressions consistent with established lineage trajectories. Held-out time-point projections further supported coarse temporal generalization, suggesting that embeddings can encode continuous biological transitions and support analyses of dynamic processes such as differentiation.

Integrating gene expression data and literature through a shared text modality addresses a limitation in current single-cell foundation models. Previous approaches that use language models to embed gene identities or metadata [2, 7] often do not incorporate the rich contextual knowledge in the biomedical literature. By leveraging this information, our approach enables interpretability and hypothesis generation across biological layers, supporting downstream tasks such as context-aware cell type annotation and the transfer of functional and disease-related knowledge into single-cell analyses. Our current implementation relies on relatively lightweight models (110M parameters), which can be efficiently run and fine-tuned even with limited hardware and are adaptable to new datasets with known cell types. Although our framework is based on compact encoder-only models, the results illustrate more broadly that adding contextual annotations represents a promising application for language models, including also large-scale language models. Our results indicate that existing general-purpose generative language models already capture aspects of biological metadata, such as relationships between cell types and their functionality, suggesting that further exposure to gene expression data could be a promising strategy to improve their applicability to genomic data analysis. Rather than a replacement for either classical scRNA-seq analysis or large generalist foundation models, our framework provides a dataset-adapted contextualization strategy. Compared with dedicated annotation or integration tools, its main purpose is not to optimize a single benchmark, but to provide a shared representation in which cells can be enriched with several layers of contextual information, such as textual labels, functional descriptions, disease-related concepts, and developmental patterns.

Nonetheless, the current implementation has important limitations. The approach relies on available cell type annotations and metadata, and is therefore semi-supervised rather than fully unsupervised. Literature retrieval may omit details found only in full-text articles and may under-represent newly described, rare, dataset-specific or inconsistently named cell states, introducing biases toward well-studied cell types, canonical functions, and frequently used terminology. We therefore report retrieval counts as a potential diagnostic to help interpret literature-informed contextualization (Additional file 2, Tables R1-R4), particularly in analyses where literature-derived text contributes strongly, such as annotation with functional descriptions. Further, employing cell sentences for converting scRNA-seq data into a text-based format for use in a language model-based framework is a pragmatic choice to allow for flexible integration with other text-based information such as metadata and literature. Yet, cell sentences also discard quantitative information from the expression matrix, although sentence-length sensitivity analyses provide an initial assessment of this loss (Additional file 1, Fig. S1). Future extensions could incorporate expression bins, rank-based encodings, or other tokenization schemes that preserve more quantitative information while retaining compatibility with text-based contextualization.

We have focused on training specialist models, designed for targeted integration of domain-specific knowledge into expression data, where prior labels or context are available. In contrast to transcriptome-text models such as CellWhisperer [18], which align quantitative transcriptomic embeddings with text representations of metadata, our framework represents all information sources in text form and uses dataset-specific literature for fine-tuning a lightweight model. At the same time, large-scale approaches like CellWhisperer and C2S-Scale [10] show the potential of generalist models trained on large datasets. Training larger base models on broader training datasets and literature corpora with our proposed training strategy could enable robust zero-shot generalization across more diverse tissues and organisms.

Additionally, future work should incorporate additional data sources such as full-text articles, curated gene and protein databases, ontologies, and pathway annotations. As single-cell datasets continue to grow in scale and complexity, integrating them with such diverse knowledge sources will be an essential step towards transitioning from task-specific models to models applicable across a broader range of biological contexts.

## 4 Conclusion

We demonstrate the utility of encoder-only language models as lightweight tools for enriching singlecell RNA-seq data with contextual information from biomedical literature. Our approach shows how expression profiles and literature-derived knowledge can be aligned in a joint embedding space and enables interpretable analyses across multiple biological layers such as cell identity, function, disease association, and developmental stage across biomedical contexts. This strategy offers a scalable and efficient framework for enriching single-cell analyses, using language models as complementary analysis tool rather than placing them at the core of single-cell data analysis. This work thus contributes to ongoing efforts to adapt language model-based representations to complex biological datasets, enabling single-cell analyses that are more directly connected to biomedical knowledge.

## 5 Methods

### 5.1 Preprocessing of single-cell RNA seq data

#### 5.1.1 Human Immune Health Atlas (Allen Institute)

We used a subset of the Human Immune Health Atlas from the Allen Institute (HIAI) containing around 230,000 T cells [34]. Cell-type annotations are provided at three levels of resolution; we used the intermediate annotation, resulting in 10 distinct T cell subtypes. We preprocessed the data using scanpy, according to the workflows described in Heumos, Schaar, et al. [26]. Cells were retained if they expressed more than 500 and less than 5000 genes, and genes were retained if detected in at least 50 cells. After normalization and log transformation of the counts, we selected the top 2000 highly variable genes (seurat_v3 method), excluding ribosomal and mitochondrial genes.

Data were split at the donor level into training (50%) and test (50%) sets, such that cells from donors used for evaluation were not included during model training or validation. This resulted in around 115,000 cells in both training and test dataset, with the training set further divided into a training (90%) and validation (10%) set. The same held-out donor split was used across all ablation models and annotation benchmarks (see Additional file 2, Fig. R1, Table R1 for an overview of the train-test design with held-out donors). Available metadata included cell type annotation, donor identity, and cytomegalovirus (CMV) status. CMV status was used only for evaluation of disease-associated shifts and was not included in cell sentences for the CMV analysis. We refer to this dataset as the *HIAI dataset*.

#### 5.1.2 Developing Mouse Brain dataset (LaManno et al.)

As a second dataset, we used the developing mouse brain atlas from La Manno, Gyllborg, et al. [43], containing approximately 270,000 cells sampled across 20 embryonic days E7-E18. We only included cell types represented by more than three cells, resulting in 130 of the 133 original cell types. We removed cells with less than 100 expressed genes and genes expressed in less than five cells. We performed library-size normalization and log transformation and selected the top 2000 highly variable genes using the seurat_v3 method, excluding ribosomal and mitochondrial genes.

For developmental analyses, we split the data into a training and test (10%) set, with the training set subdivided into a training (90%) and validation (10%) set, so that the evaluation was performed on held-out mice. In an additional supplementary analysis, selected embryonic time points were withheld during training and cells from these time points were projected into the learned embedding space after model fitting (see Additional file 1, Fig. S10, Additional file 2, Fig. R2). We refer to this dataset as the *LaManno dataset*.

#### 5.1.3 Small PBMC dataset for sentence-length and seed sensitivity analyses

To evaluate the effect of cell sentence length and random seed on a computationally tractable example, we used a smaller peripheral blood mononuclear cell (PBMC) dataset. The dataset was obtained from the 10X Genomics database [46]. We refer to it as the *PBMC* dataset. The dataset was processed analogously to the HIAI dataset, including normalization, log transformation, highly variable gene selection, and construction of cell sentences with different numbers of top-ranked genes. This analysis was used only as a supplementary sensitivity analysis and was not used to tune the larger HIAI or LaManno analyses.

### 5.2 Generation of cell sentences

To represent high-dimensional quantitative gene expression profiles in textual form, we followed the Cell2Sentence approach [14]. For each cell, we ranked highly variable genes (excluding housekeeping genes) by expression, and concatenated the top-ranked gene symbols into a gene list. Unless stated otherwise, downstream analyses used cell sentences containing the top 50 ranked genes.

We evaluated the effect of sentence length in a supplementary analysis on the PBMC dataset. Cell sentences were generated with lengths ranging from 10 to 200 genes. To quantify how much expression-derived structure was retained, we transformed each sentence representation into a binary gene-incidence matrix indicating whether a gene was present in the sentence (this transformation discards the rank-based order of the genes, thus resulting in some underestimation of the variation). We then applied principal component analysis (PCA) [47] to this binary matrix, and compared the number of principal components needed to explain 90% of the variance with the corresponding value for the log-normalized gene expression matrix restricted to 2000 highly variable genes. We also recorded the total number of unique genes represented across cell sentences for each sentence length. In addition, models trained with different sentence lengths (10, 20, 50, 100, 200) were evaluated for mean cell type label-alignment receiver operating characteristic area under the curve (ROC-AUC) and mean biological conservation score from [31] across three random seeds (Additional file 1, Fig. S1). These analyses supported the use of 50 genes as a pragmatic default that retained expression-derived structure while keeping sentence length computationally manageable. In all analyses shown in the manuscript, we therefore used ranked gene lists truncated at a length of 50.

For the training data, we then used sentence templates to convert gene lists into “semantic” cell sentences. Templates either contained only the ranked gene list or combined the gene list with available metadata and included examples such as:

- Gene list only: “*{gene_str} are the top expressed genes in this cell.*”
- Gene list and cell type: “*A cell that expresses the following genes: {gene_str} is likely a {celltype} cell.*”
- Gene list, cell type and time point: “*A {celltype} cell at {time} expresses these genes: {gene_str}.*”

For training with different metadata information, we combined cell sentences of gene lists only with sentences including also metadata. For example, in experiments combining gene lists and cell type information, we used sentences containing only the gene list for 30% of cells, and sentences containing both the gene list and the cell type label for 70% of cells. In experiments combining gene lists, cell type and time metadata, sentences were distributed across four variants: sentences containing the gene list, the cell type label and the time point (50%), sentences containing the gene list and time point (20%), sentences containing the gene list and cell type (20%), and sentences containing the gene list only (10%).

All downstream evaluations used metadata-free cell sentences. Specifically, test data were converted to ranked gene lists without appended metadata, such that evaluation cell sentences did not contain cell type labels, donor metadata, CMV status, or embryonic day. This metadata-free evaluation was used for held-out donors in the HIAI analyses and held-out mice in the developmental analyses. For the PBMC dataset a random stratified test split was generated.

For shuffled-label controls, labels were randomly permuted within the corresponding training data before triplet generation. Because shuffling was implemented as a permutation of the existing label vector, the marginal label distribution was preserved exactly. Seeds are reported in the scripts and configuration files in our GitHub repository (https://github.com/MariaKrissmer/alias).

### 5.3 Collecting data for training on literature

For enriching scRNA-seq datasets with additional information from scientific literature, we collected titles and abstract relevant to the cell types within the datasets from PubMed records and curated cell type knowledge sources. PubMed titles and abstracts were retrieved through the NCBI Entrez interface [21]. Searches were performed separately for each dataset and were targeted to the organism, cell types, and, where applicable, metadata of interest represented in the corresponding scRNA-seq dataset. Search terms included (1) the organism as a MeSH term (human or mouse), (2) the cell types of interest, i.e., the ones present in the considered dataset, and (3) optional additional metadata of interest, such as embryonic day. Queries used Boolean operators to combine or exclude metadata-specific terms.

Example query templates were (see Additional file 2, Tables R2-R4 for a full list):

- Cell type query: homo sapiens[MeSH] AND Blood Cells[MeSH] AND “{celltype}”[Title/Abstract]
- Cell type and disease query: homo sapiens[MeSH] AND Blood Cells[MeSH] AND “{celltype}”[Title/Abstract] AND {disease}
- Cell type excluding disease query: homo sapiens[MeSH] AND Blood Cells[MeSH] AND “{celltype}”[Title/Abstract] NOT {disease}

For the HIAI T-cell literature datasets, we retrieved up to 3000 PubMed records per cell type query. Titles and abstracts were processed separately and split into token-limited segments using the PubMedBERT tokenizer. Each text segment was assigned a label based on the query from which it was retrieved. Records associated with multiple labels were removed where configured to avoid ambiguous supervision.

In addition to PubMed text, we appended curated Cell Ontology [28] and ScType-derived [29] text entries for the HIAI dataset only. Cell Ontology entries were generated from a curated mapping between the cell type labels and Cell Ontology (CL) terms. We included CL definitions and description sentences for mapped cell types, split long descriptions into individual sentences, and excluded sentences shorter than eight words. For held-out-cell-type literature variants, entries corresponding to held-out labels were excluded. ScType marker information was added as structured text statements listing positive marker genes for each cell type. These marker rows were treated as cell-type-associated knowledge entries and assigned the corresponding cell type label.

The resulting literature pool for the therefore contained three text sources: PubMed titles/abstracts, Cell Ontology definitions/descriptions, and ScType positive marker-gene statements. For HIAI T-cell literature-based dataset generation, Cell Ontology and ScType rows were converted into title-like and abstract-like training rows so that they entered the same triplet-generation pipeline as PubMed-derived text. For the CMV analysis, we trained one model with CMV-related literature queries and one control model without dedicated CMV-specific literature queries. In both settings, CMV metadata was withheld from cell sentences. The control model was used to assess whether the observed CMV-associated cytotoxicity signal depended on dedicated CMV-specific literature or was also represented in broader cell-type-associated literature.

For each literature query, we recorded the number of retrieved titles, abstracts, and database entries. Query strings, retrieval counts, and retrieved record identifiers are reported in Additional file 2, Tables R2-R4 and in the code repository with the corresponding datasets retrieval scripts respectively.

### 5.4 Base model

All sentence embedding and fine-tuning tasks of our approach were initialized from PubMedBERT (neuml/pubmedbert-base-embeddings, available via Hugging Face) as base model, a domain-specific Bidirectional Encoder Representations from Transformers (BERT) model trained on Pubmed titles and abstracts [15, 48]. The base PubMedBERT model is a 12-layer bidirectional Transformer encoder with 110M parameters, and outputs 768-dimensional embeddings.

The original PubMedBERT model has been trained on a large-scale corpus of parsed and indexed PudMed titles and abstracts (via the paperetl pipeline), with training pairs created from combinations of titles and corresponding abstracts or semantically similar titles. Training used the Multiple Negatives Ranking Loss (for details, see below in Section 5.9) with mean pooling for token aggregation, which encourages semantically similar pairs to be closer in embedding space. PubMedBERT outperforms general-purpose embedding models on biomedical semantic similarity tasks (e. g., PubMedQA, PubMed Subset, PubMed Summary).

This pre-trained encoder served as the backbone for all model fine-tuning experiments in this study.

### 5.5 Siamese Sentence BERT architecture

To enable efficient similarity computations, we used the pre-trained PubMedBERT model within a Sentence-BERT (SBERT) framework [16], a modification of BERT designed for efficient semantic similarity in sentence-pair tasks, such as aligning cell sentences of similar cells. In contrast to standard BERT, SBERT encodes each sentence independently using a siamese or triplet architecture [49]. A mean pooling layer produces a fixed-size (768-dimensional) token embedding, which enables fast pairwise similarity computation using metrics such as cosine similarity. This architecture reduces the number of model passes required for pairwise comparisons [16], making it well suited for large-scale comparison of cell sentences and literature-derived information.

SBERT supports training with contrastive objectives such as triplet loss functions, contrastive loss functions, and multiple negatives ranking (MNR) loss functions, which we employed in our training procedure (see Section 5.9 for details). Because the model is designed to cluster semantically similar sentences in embedding space, it is well suited for tasks like semantic alignment and clustering.

All downstream and fine-tuning analyses in this work are therefore based on a PubMedBERT encoder within an SBERT architecture.

### 5.6 Generation of triplets for contrastive fine-tuning using label-aware random-positive and label-aware hard-negative-mining

For each dataset, we constructed label-aware triplets for fine-tuning the PubMedBERT sentence embedding base model. Each triplet consisted of an anchor sentence, a positive sentence sharing the same label, and a hard negative sentence with a different label but high embedding similarity. Unless stated otherwise, we sampled *J* = 5 positive and negative examples per anchor. With these triplets, we used the multiple negatives ranking (MNR) loss as fine-tuning objective, to generate embeddings which capture meaningful semantic interactions of genes, cells, cell types and additional information such as embryonic time.

#### 5.6.1 Triplet generation for scRNA-seq data

We use each semantic cell sentence *x* ∈ {*x*_1_*,…,x_N_*} as anchor (see Section 5.2) and sample positive and negative cell sentences to construct *J* = 5 triplets per anchor cell. Specifically, for an anchor cell sentence *x*, we sampled *J* positive examples *x*^+^*, j* = 1*,…,J* from other cells sharing the same cell type label. Negatives were selected using label-aware hard negative mining: for an anchor *x*, we identified the *J* most similar sentences with different cell type labels, where similarity was measured using cosine similarity in the PubMedBERT base embedding space (see Section 5.9). Each anchor-positive pair (*x, x_j_*^+^) for a *j* ∈ {1*,…,J*} was paired with one unique hard negative *x_j_*^-^ for a *j* ∈ {1*,…,J*}, resulting in *J* anchor-positive-negative triplets per anchor cell. Formally, the resulting training set is given by

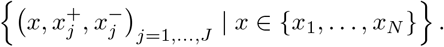

This approach avoids the quadratic expansion of standard hard negative mining, where each anchor-positive pair is assigned *k* negatives, resulting in *k*^2^ triplets, while maintaining diversity. We implemented hard negative mining using a modified version of the corresponding function implemented within the sentence-transformers package to account for cell type labels and generate triplets with unique negatives to avoid dataset explosion. The label-aware hard negative mining was run in batches of 10,000 cells.

#### 5.6.2 Triplet generation for literature-derived text

Titles, abstracts, and database entries retrieved from literature sources were treated analogously to cell sentences. Each title served as an anchor with five positive randomly sampled from titles with the same label, where labels were derived from the search query. Positive examples were sampled from texts retrieved with the same query-derived label, and hard negatives were selected from texts with different labels based on cosine similarity in the base embedding space. For PubMed abstracts, each anchor abstract was randomly assigned positive abstracts returned by the same search query. Negatives for titles, abstracts, and database entries were obtained by label-aware hard negative mining as described for scRNA-seq-derived cell sentences. All triplets from both scRNA-seq-derived and literature-derived data were combined for model fine-tuning.

### 5.7 Ablation model design

To assess how different information sources contribute to the learned representation, we trained matched ablation models that varied the inclusion of scRNA-seq-derived cell sentences, metadata-containing cell sentences, literature-derived text, and shuffled controls. All ablation models were initialized from the same base model and trained with the same fine-tuning hyperparameters unless stated otherwise. The following model variants were used:

- MA: full model trained on scRNA-seq-derived cell sentences, cell type labels/metadata, and literature-derived text.
- MA*: full model trained as MA, but with shuffled labels for literature-derived text.
- MB: model trained on scRNA-seq-derived semantic cell sentences with cell type labels/metadata, without literature-derived text.
- MC: model trained on cell sentences without appended cell type labels/metadata, using the correct cell type labels for triplet construction.
- MC*: model trained on cell sentences without appended cell type labels/metadata, using shuffled cell type labels for triplet construction.
- MD: model trained only on literature-derived text.
- Base: pre-trained sentence embedding model without additional fine-tuning.

For shuffled controls, labels were randomly permuted while preserving the marginal label distribution. In MA*, literature labels were shuffled before literature triplet construction. In MC*, cell type labels were shuffled before scRNA-seq triplet construction. These controls preserve the amount and format of the input text while disrupting the correspondence between biological labels and training pairs.

### 5.8 Fine-tuning of models using the sentence-transformer package

We fine-tuned sentence embedding models using the sentence-transformers package. For joint models, fine-tuning alternated epoch-wise between scRNA-seq-derived cell sentence triplets and literature-derived triplets to expose the model to both data sources while avoiding direct mixing of batches from heterogeneous text sources. Literature-only and cell-only ablation models used the corresponding subset of triplets.

Training was performed with a batch size of 128 for both literature-derived and scRNA-seq-derived data. Models were trained for 15 epochs with 1000 warm-up steps, weight decay of 0.01, mixed precision training (fp16=True), and gradient clipping with max_grad_norm=1.0, the best checkpoint was selected using a joint validation score on both evaluation splits where applicable. Random seeds for data splitting, triplet sampling, label shuffling, and model training are included in the metadata files for all models together with the corresponding training scripts in our code repository. For ablation study models only trained on either an scRNA-seq-based or a literature-based dataset we trained for 8 epochs, while keeping all other training settings equivalent.

Because of limited compute resources and the number of individual models and evaluations used in the manuscript, most large-scale analyses were run once per model variant. To evaluate stochastic variability in a computationally tractable setting, the PBMC sentence-length sensitivity analysis was repeated across three random seeds (Additional file 1, Fig. S1). Models were trained for 8 epochs on the scRNA-seq derived datasets only, using a batchsize of 64 across all cell sentence lengths.

### 5.9 Multiple negatives ranking loss

We fine-tuned all models using the multiple negatives ranking (MNR) loss function [24], a contrastive objective related to the InfoNCE loss. Given a batch of size *M < JN* of anchor-positive pairs, the loss increases a similarity measure *S_c_* between an anchor *x* and its positive paired sample *x_j_*^+^*, j* ∈ {1*,…,J*}. As similarity measure we use cosine similarity, defined by

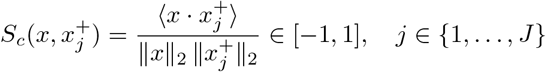

The MNR loss function treats all other positives in the batch *M* as implicit negatives. For a given anchor-positive pair (*x, x*^+^), where *j* ∈ {1*,…,J*}, the standard MNR loss is:

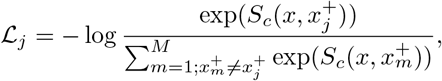

and all positives *x_m_*^+^ for *m* ≠ *j* are treated as in-batch negatives.

From a probabilistic interpretation, the loss function maximizes the log-likelihood of correctly identifying the true positive, where the softmax over all candidates ensures that the similarities are normalized to a probability distribution, i. e., it resembles a softmax classifier over cosine similarities. It is minimized when the similarity between an anchor and its true positive is much larger than to any of the in-batch negatives.

To improve generalization, we combined these soft in-batch negatives with label-aware hard negatives obtained as described above (see Section 5.6). Given a triplet of anchor-positive-negative pairs (*x, x_j_*^+^,*x_j_*^-^)*_j_*_=1_*_,…J_*, obtained as described above in Section 5.6.1, we extend the loss for one anchor-positive pair (*x, x_j_*^+^) for *j* ∈ {1*,…,J*} to:

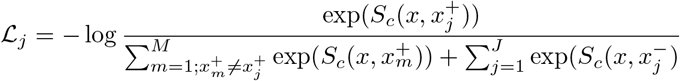

Each anchor–positive pair was therefore trained against both random in-batch negatives and carefully selected near-miss examples. This encourages the model to separate closely related but differently labeled biological examples while aligning semantically related cell-derived and literature-derived texts and increase robustness to noise. Further, the exposure to a broader range of negative samples helps reduce overfitting.

### 5.10 Cosine similarity of cells with cell type labels

We computed the cosine similarity *S_c_* between held-out cell embeddings *c*_1_*,…,c_N_* and embeddings of candidate cell type labels *l*_1_*,…,l_L_*:

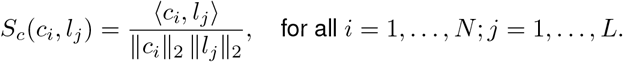

For label-alignment analyses, we compared the distribution of similarity scores between a given cell type label and cells annotated as that type with the distribution of similarity scores between the same label and all other cells. We used one-vs-rest ROC curves to quantify how well similarity scores separated cells of each type from all other cells and reported the area under the ROC curve (ROCAUC) [50] for each cell type.

For multiclass cell type annotation, each held-out cell was assigned to the candidate cell type label with the highest cosine similarity. Annotation performance was evaluated using balanced accuracy, macro F1 score, and weighted F1 score.

For synonym-based evaluation, candidate cell type labels were expanded using synonyms derived from the curated HIAI-to-Cell Ontology mapping from the literature-based dataset generation (see Methods Section 5.3). For each reference cell type, we included the canonical dataset label together with available Cell Ontology (CL) labels, synonyms, and exact synonyms from the matched CL term. Case-only variants and duplicate synonym strings were removed. Each synonym was embedded and used as an alternative query label, and predictions made to synonym labels were mapped back to the corresponding canonical cell type reference label before computing annotation metrics. We used this to evaluate whether annotation performance was robust to biologically equivalent terminology rather than limited to matching an exact dataset label.

### 5.11 Comparison with CellTypist and SingleR

To provide reference points for cell type annotation performance, we compared our similarity-based annotation strategy with CellTypist [37, 38] and SingleR [39]. Both methods were trained or configured using the same held-out donor split as the embedding models. Training donors were used as reference data, and performance was evaluated on cells from held-out donors.

For CellTypist, we trained a dataset-adapted local model on the reconstructed HIAI T-cell training split. Training used the raw count representation and the cell type labels used for triplet generation and evaluation in our models. The model was trained with celltypist.train using the default training parameters. The trained model was saved as a local CellTypist model and then applied to held-out donor cells with celltypist.annotate.

For SingleR, we used the same reconstructed HIAI T-cell training split as the reference dataset and the same held-out donor dataset as the query. SingleR was run with the Python singler.annotate_single interface. Query and reference matrices were restricted to their shared genes, represented as feature-by-cell matrices, and the same reference labels as for all other analyses were used. The reference was capped at at most 1,000 cells per label using a fixed sampling seed, and annotation was run with default SingleR training and classification arguments.

For both baselines, we standardized predictions to the same output format as the embedding-based annotation results. We computed balanced accuracy, macro F1 score, and weighted F1 score against the HIAI reference annotation and compared them with the corresponding metrics from our similarity-based annotation approach (Additional file 1, Fig. S3 **A**, **B**).

### 5.12 Embedding quality benchmark

We evaluated embedding quality using single-cell integration metrics for biological conservation, batch correction, and total score following the scIB benchmark framework [31]. Biological conservation and batch correction metrics used the default BioConservation() and BatchCorrection() metric sets from scib-metrics. The total score was the default scIB aggregate, corresponding to 0.6 times biological conservation plus 0.4 times batch correction.

All embeddings were evaluated on the same held-out HIAI T-cell donor test split. If more than 20,000 test cells were available, we subsampled cells. Embeddings from the full model and ablations were generated from the held-out test cell sentences with batch size 256 and compared with single-cell representation baselines. Specifically, as embedding baselines we included three PCA-based representations, scGPT embeddings [2], and scVI embeddings [32] based on models pre-trained on the CELLxGENE Census [33].

We computed a baseline PCA representation with 50 components on the processed gene expression data. Additionally, we computed two PCA versions based on ranked cell sentence gene lists: PCA_rank_weighted used sparse cell-gene values weighted by inverse square-root rank, and PCA_one_hot used binary presence or absence of a gene in the cell sentence.

scGPT embeddings were generated with the local scGPT checkpoint specified for the run following the scGPT zero-shot embedding workflow. As input we used the held-out test data counts with gene names, the maximum sequence length was 1,200 genes with batch size 64.

For scVI, we generated two CELLxGENE Census baselines. The pretrained baseline used the Census scVI model for homo_sapiens from Census version 2025-11-08 and performed encoderonly inference without query adaptation. As a dataset-adapted baseline, we used the same reference model but performed scVI query adaptation on the HIAI test data before extracting latent representations. Query adaptation used the gene expression counts with a batch size of 1,024 and standard training parameters.

We evaluated all embeddings on the same held-out HIAI donor cells and the same cell type and donor annotations. Full metric tables are reported in Additional file 1, Fig. S1.

### 5.13 Evaluation of contextualization with functionality descriptions

#### 5.13.1 Expert-curated functionality descriptions

To evaluate whether embeddings captured functional information beyond cell type labels, we curated expert-defined functionality descriptions for T cell subtypes in the HIAI dataset. Descriptions, corresponding expert-assigned cell types, and supporting references or rationale [51, 52, 53, 54, 55] are provided in Additional file 1, Figure S4. We additionally generated paraphrased versions for the MAIT cell functionality description to assess robustness to rewording.

We evaluated functionality descriptions at two levels. At the cell type label level, we embedded each functionality description and each candidate cell type label, and ranked candidate cell types by cosine similarity to the functionality description.

At the single-cell level, we computed similarity scores between the embeddings of each held-out cell sentence and each functionality description. For each description and candidate cell type, we compared the distributions of the similarity scores for cells of the respective cell type with the scores of all other cells. By varying the decision threshold over the similarity scores, we generated ROC curves that quantified how well cell-wise similarity scores separated cells of the candidate type from all other cells based on their similarity scores to the given functionality description. The resulting area under the ROC curve thus provides a measure of the association between a functionality description and each cell type, reflecting how distinctively the description matches the cells’ embeddings. We thus summarized the cell-level association by computing these one-vs-rest ROC-AUC values, and ranked cell types by this cell-level association score.

For each functionality description *i*, we recorded the rank *r_i_* of the expert-assigned cell type and summarized performance across descriptions using mean reciprocal rank (MRR):

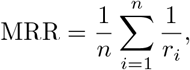

where *n* is the number of functionality descriptions. An MRR of 1 indicates that the expert-assigned cell type was ranked first for every description.

#### 5.13.2 Comparison with CellWhisperer and generative language models

For comparison with CellWhisperer [18], we used the same expert-curated functionality descriptions and the same cell-level evaluation strategy. To generate the embeddings of the single cells of the HIAI data and the query texts, we loaded the checkpoints of the published CellWhisperer model (https://medical-epigenomics.org/papers/schaefer2025cellwhisperer/data/models/cellwhisperer_clip_v1.ckpt) and embedded raw counts using the provided helper functions adata_to_embeds() and score_transcriptomes_vs_texts(). For scoring, we used the logit scale of the loaded model and a softmax normalization. All analyses where conducted with the mmcontext framework [56].

For comparing our approach to general-purpose generative language models, we used the celltype-based comparison strategy. Models were prompted to choose which of two candidate T cell types more likely matched a given functionality description. We queried Llama3.3 70B [57] and Qwen3 235B [58] via OpenRouter [59]. The prompt was:

*You are a biomedical expert specializing in T cell immunology.*

*Functional description: {description}*

*Option A: {celltype_a}*

*Option B: {celltype_b}*

*Question: Which of these two T cell types most likely performs the described function? Please respond ONLY with one of the following exactly:*

*A*

*B*

*Do not add any other text*.

For each functionality description, all possible ordered pairs of candidate cell type labels were presented to the model, i.e., each pair was presented in both possible orders to eliminate potential position bias in the model’s responses. Each model response was recorded as a “win” for the selected cell type. Across all pairs, the total number of wins per cell type was normalized by the maximum attainable number of wins and converted into a rank, and the rank of the expert-assigned cell type was used for comparison with the embedding-based methods.

#### 5.13.3 Cell-type-overarching functional queries

To evaluate functional programs that span multiple cell types, we embedded short functional queries and computed cosine similarity between each held-out cell embedding and the query embedding. We focused on the queries “cytotoxic” and “regulatory”. The cytotoxicity query was used in the main analysis, and analogous analyses for the regulatory query are provided in Additional file 1, Fig. S5.

#### 5.13.4 Automatic thresholding and sensitivity analyses

For each functional query, we used Otsu’s method [40] to determine an automatic threshold on the distribution of cell-wise similarity scores. Cells with similarity scores above the threshold were assigned to the high-similarity group for the corresponding functional query, and cells below the threshold were assigned to the low-similarity group.

To evaluate whether downstream results depended on the exact cutoff, we performed threshold sensitivity analyses around the selected Otsu threshold. For each query, thresholds were sampled across the observed similarity-score range, starting from the Otsu threshold plus or minus 10% of the score range and bounded by the observed minimum and maximum scores. For each threshold, we reassigned cells to high-and low-similarity groups and recorded the number and fraction of high-similarity cells and the mean gene-set score in each group. Differential expression was performed for the Otsu-based assignment using Wilcoxon rank-sum implementation from scanpy. We then tracked the top differentially expressed genes from this selected cutoff across the sensitivity thresholds by recomputing their mean expression difference between high-and low-similarity cells. Cytotoxicity marker recovery was evaluated for the main cytotoxicity query, and the same workflow was repeated for the regulatory query in the supplementary material.

#### 5.13.5 Comparison with gene-set scores

We compared embedding-based functional similarity scores with conventional gene-set scores. Gene-set scores were computed using the scanpy.tl.score_genes function with curated cytotoxicity (*NKG7*,*GZMA*, *CTSW*, *CST7*, *CCL5*) and regulatory (*FOXP3*, *IL2RA*, *CTLA4*, *IKZF2*, *TNFRSF18*, *TIGIT*) gene sets.

For each query, cells were ranked separately by embedding-based similarity score and by gene-set score. We divided each ranking into deciles and computed a decile-by-decile concordance table summarizing how cells ranked by one score were distributed across deciles of the other score.

We additionally evaluated recovery of top-ranked gene-set-score cells among cells with high embedding-based similarity. Specifically, we compared the overlap between the top percentiles of cells ranked by gene-set score and the top percentiles of cells ranked by embedding similarity. These analyses were used to assess whether cells most strongly associated with a functional program by conventional marker-based scoring were also prioritized by the text-based embedding query.

### 5.14 Disease-associated functional analysis

To evaluate disease-associated functional shifts, we focused on CMV-associated cytotoxicity in the HIAI T-cell dataset. CMV status was not included in cell sentences. Evaluation was performed on held-out donors using cell sentences without metadata. We embedded the disease-associated query “increased cytotoxic activity in cytomegalovirus positive patients” and computed cosine similarity between the query embedding and the embeddings of each held-out cell.

We employed the cell-based comparison level strategy described in Section 5.13 for assessing the similarity of single cells to a disease interaction description. Within each cell type, we compared the distribution of cell-wise similarity scores between cells from CMV-positive and CMV-negative donors using a Mann-Whitney U test. As cells from the same donor are not statistically independent, cell-level tests are used as descriptive comparisons and not interpreted as donor-level inferential evidence. Further, we compared two groupings for differential expression analysis for memory CD4+ T cells. First, cells were grouped by donor-level CMV metadata. Second, we grouped cells by model-derived similarity to the CMV-associated query and used Youden’s J [42] to determine the optimal similarity score threshold for separating cells from CMV-positive and CMV-negative individuals. We did not adjust for donor-level structure and accordingly treated *p*-values as descriptive. We interpreted these results as exploratory plausibility checks and as a means for relative cell-level comparisons only.

A supplementary control model trained without dedicated CMV-specific literature queries was evaluated with the same procedure.

### 5.15 Differential expression analyses

Differential expression analyses were performed with scanpy v1.11.5 using scanpy.tl.rank_genes_groups. Cells were grouped by model-derived similarity to given embedding strings, and genes enriched in the high-similarity group were tested against the low-similarity reference group using a Mann-Whitney U test (method =’wilcoxon’). As input values, we used normalized and log-transformed gene expression counts from held-out cells. *P*-values, adjusted *p*-values, and log-fold changes were taken from the scanpy results table, using the default multiple-testing correction (Benjamini-Hochberg). For visualization, adjusted *p*-values were plotted as − log_10_(adjusted *p*-value).

For functional query analyses, differential expression compared cells above and below the functional similarity threshold for each tested cutoff. For CMV analyses, differential expression was restricted to memory CD4+ T cells and compared groups defined either by donor-level CMV metadata or by model-derived CMV-associated similarity. These analyses were used as descriptive plausibility checks for recovery of known marker genes and for comparing model-derived and metadata-derived groupings rather than treated as donor-level inference and not intended as de novo gene signature discovery.

### 5.16 Developmental trajectory and pseudotime analyses

For developmental analyses, we evaluated embeddings on held-out mice using cell sentences that did not contain cell type labels or embryonic day metadata. To visualize broad developmental organization, the LaManno dataset was divided into seven subsets (see Additional file 1, Fig. S7/8). UMAP embeddings were computed from model embeddings and colored by reference cell type annotation and embryonic day.

For the forebrain trajectory analysis, we selected cells annotated as ‘Dorsal forebrain’, ‘Forebrain’, ‘Ventral forebrain’, ‘Neuronal intermediate progenitor’, ‘Forebrain glutamatergic’, ‘Forebrain GABAergic’, and ‘Cortical or hippocampal glutamatergic’. We computed diffusion pseudotime [60] twice: once based on the model embeddings and once based on the processed gene expression representation after batch integration with scVI [32], to mitigate clearly visible batch effects. Root cells were selected from ‘Forebrain’ labeled cells, specifically the cell with the earliest developmental time point was used as the diffusion pseudotime root. For the model embeddings, we constructed the neighbourhood graph using the 15 nearest neighbors based on 50 principal components, while using the integrated scVI representation directly for the expression-based pseudotime calculation. We assessed pseudotime concordance between the model embedding and gene expression representation using Kendall’s *τ* [61] based on cell ranks.

We evaluated whether held-out time-point cells mapped to developmentally plausible regions by comparing local neighborhoods in the model embedding with neighborhoods in the scVI-integrated representation. Neighborhoods were computed within the configured forebrain lineage using Euclidean nearest-neighbor search with *k* = 15.

For each held-out cell, we computed the top-*k* Jaccard overlap between model and scVI neighbor sets based on the 15 nearest neighbors of each cell in the corresponding representation. In addition, we computed a rank-weighted neighborhood affinity matrix in which the neighbor at rank *r* was assigned weight 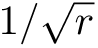 for *r* ≤ 15 and all non-neighbor cells were assigned weight zero. Weighted overlap between model embedding and gene expression neighborhoods was summarized per cell as the ratio between the summed element-wise minimum and maximum of the two rank-weighted affinity vectors. Cell-level metrics were then aggregated by embryonic time point and cell type. These analyses were used as qualitative support for coarse temporal generalization.

### 5.17 Availability of code, processed data, and model outputs

The code used for preprocessing, cell sentence generation, literature retrieval, model fine-tuning, and downstream evaluation is available at https://github.com/MariaKrissmer/alias. The repository includes configuration files documenting data splits, random seeds, triplet sampling, label shuffling, and model training settings. Query strings are provided in Additional file 2, Tables R2-R4, and PMID lists and configurations for reproducing processed cell sentences and exact train/validation/test splits are provided in the repository. Selected trained model checkpoints will be made available on Hugging Face upon publication of the manuscript.

## Declarations

### Funding

This work is funded by the Deutsche Forschungsgemeinschaft (DFG, German Research Foundation) – Project-ID 322977937 – GRK 2344 (JR, TV) and Project-ID 499552394 – SFB 1597 (SMK, HB, MH) and by the Deutsche Forschungsgemeinschaft (DFG, German Research Foundation) under Germany’s Excellence Strategy CIBSS – EXC-2189 – Project ID 390939984 (SMK, JM, HB)

### Conflict of interest / Competing interests

The authors declare that they have no competing interests.

### Consent for publication

All authors have read and approved the final manuscript and consent to its publication.

### Data availability

The study used only publicly available datasets. Specifically, we analyzed a subset of the Human Immune Health Atlas [34], restricted to T cells, and the developing mouse brain dataset from La Manno, Gyllborg, et al. [43]. Detailed preprocessing steps are provided in the Methods section.

### Materials availability

No new materials were generated in this study.

### Code availability

See Methods Section 5.17.

### Author contribution

SMK performed the research, conducted the analyses, and wrote the manuscript. JM contributed to the model implementation. JR and TV provided biological expertise and interpretation, in particular for the lineage trajectory section. HB and MH supervised and guided the work, MH co-wrote the manuscript. All authors reviewed and approved the final version.

## Notes

### Competing Interest Statement

The authors have declared no competing interest.

### Summary of Updates

This preprint has been updated to match the revised manuscript submitted following journal peer review. The revision includes ablation studies for assessment of the joint literature-enriched embedding space, improved benchmarking of annotation performance and embedding quality and all evaluations were adapted to held-out donors resulting in updated versions for all main figures. Functionality and disease-associations were expanded with sensitivity analyses and minor corrections were made throughout the manuscript. The principal conclusions remain unchanged.

