## Additional file 1 for "Adding layers of information to scRNA-seq data using pre-trained language models"

### Additional file 1 for
### Adding layers of information to scRNA-seq data using pre-trained language models

Sonia Maria Krißmer ✉[1,2], Jonatan Menger[1,2], Johan Rollin[3], Tanja Vogel[3], Harald Binder[1,2,4], Maren Hackenberg ✉[1,4,5]

[1] Institute of Medical Biometry and Statistics (IMBI), Faculty of Medicine and Medical Center, University of Freiburg, Germany
[2] CIBSS – Centre for Integrative Biological Signalling Studies, University of Freiburg, Germany
[3] Institute of Anatomy and Cell Biology, Department Molecular Embryology, Faculty of Medicine, University of Freiburg, Germany
[4] Freiburg Center for Data Analysis, Modeling and AI, University of Freiburg, Germany
[5] Present address: European Molecular Biology Laboratory, Heidelberg, Germany

August 10, 2026

### Supplementary figures

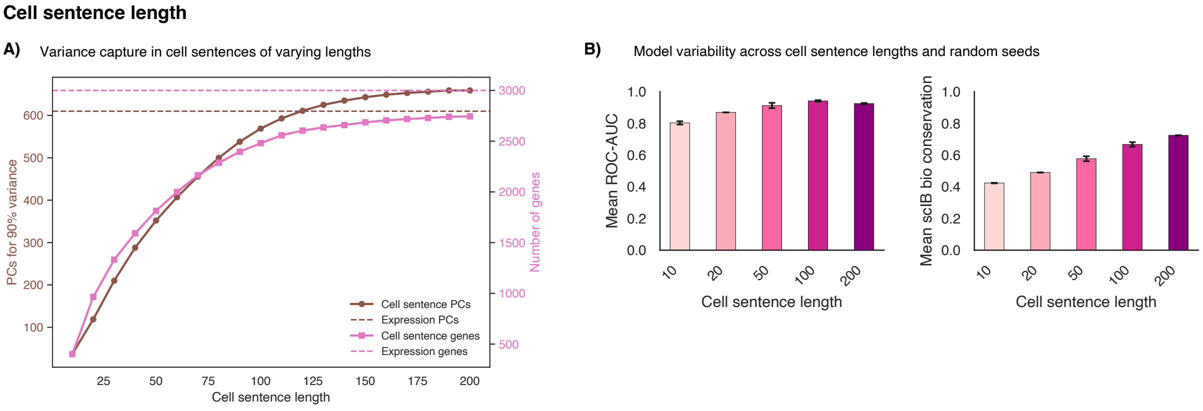


**Figure S1: Effect of cell sentence length on preserving expression structure in the embedding in a PBMC example dataset [1].** **A)** Comparison of cell sentences with different numbers of top-ranked genes. For each sentence length, cell sentences were represented as binary cell-by-gene matrices indicating whether a gene was included in the sentence, and principal component analysis (PCA) was used as a simple summary of retained structure. The number of principal components required to explain 90% of the variance is shown together with the number of unique genes represented in the corresponding cell sentences. For comparison, the same quantities are shown for the full quantitative expression matrix using 2000 highly variable genes. **B)** Mean ROC-AUC and mean biological conservation score (following the single-cell integration benchmarks from [2]) across models trained with different cell sentence lengths. Results are shown across three random seeds each.

**Marker genes for annotation of T cells**

A)

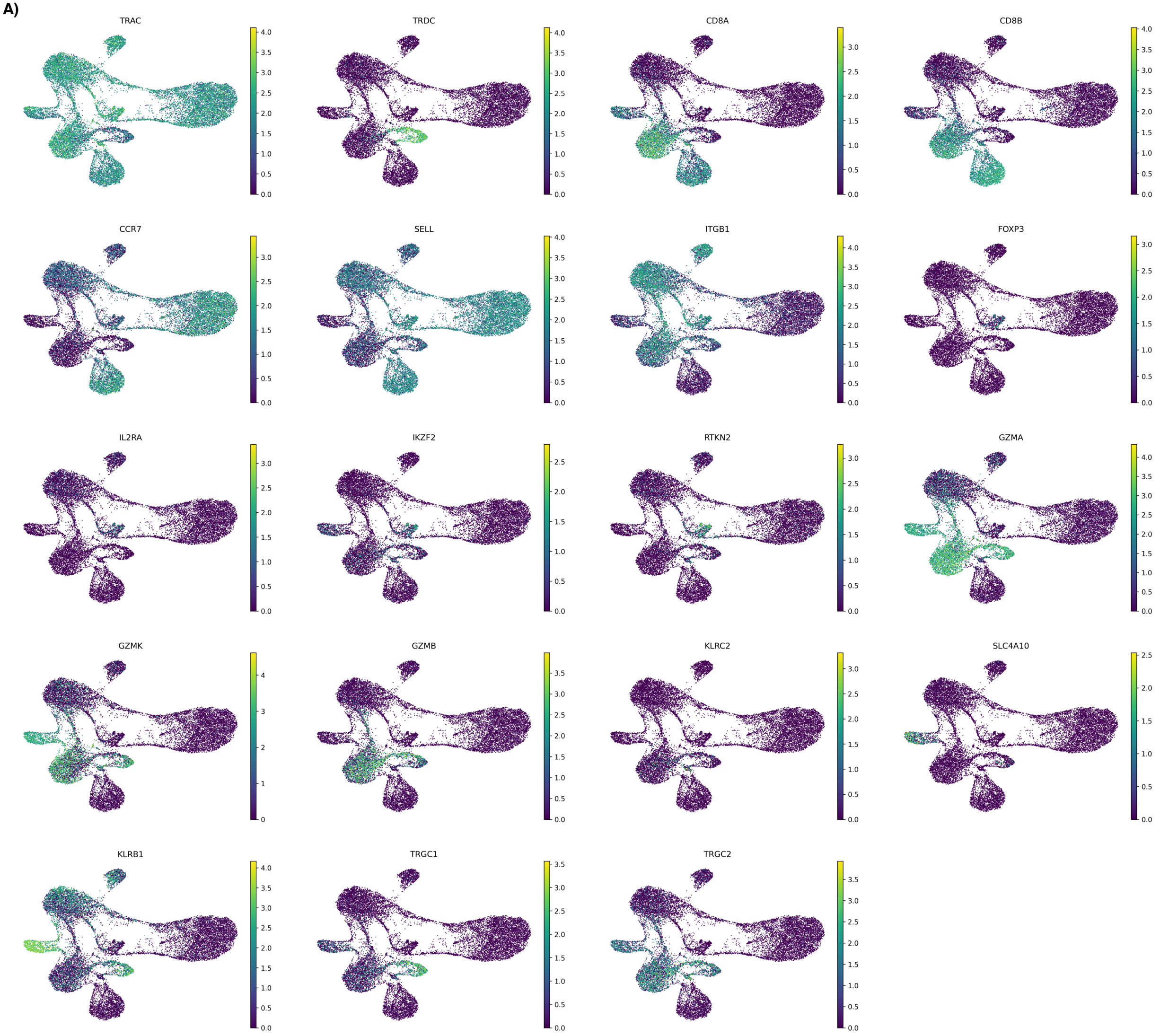


**Figure S2: Marker genes expression patterns in model embeddings for different T cell subsets in the *HIAI* dataset.** Selection of marker genes is based on the annotation strategy from the *HIAI* dataset. Expression of the selected marker genes is shown on the UMAP representation based on full-model embeddings. The marker patterns support that the learned embeddings preserve biologically meaningful T-cell subset structure and within-cell-type heterogeneity.

#### Cell type annotation

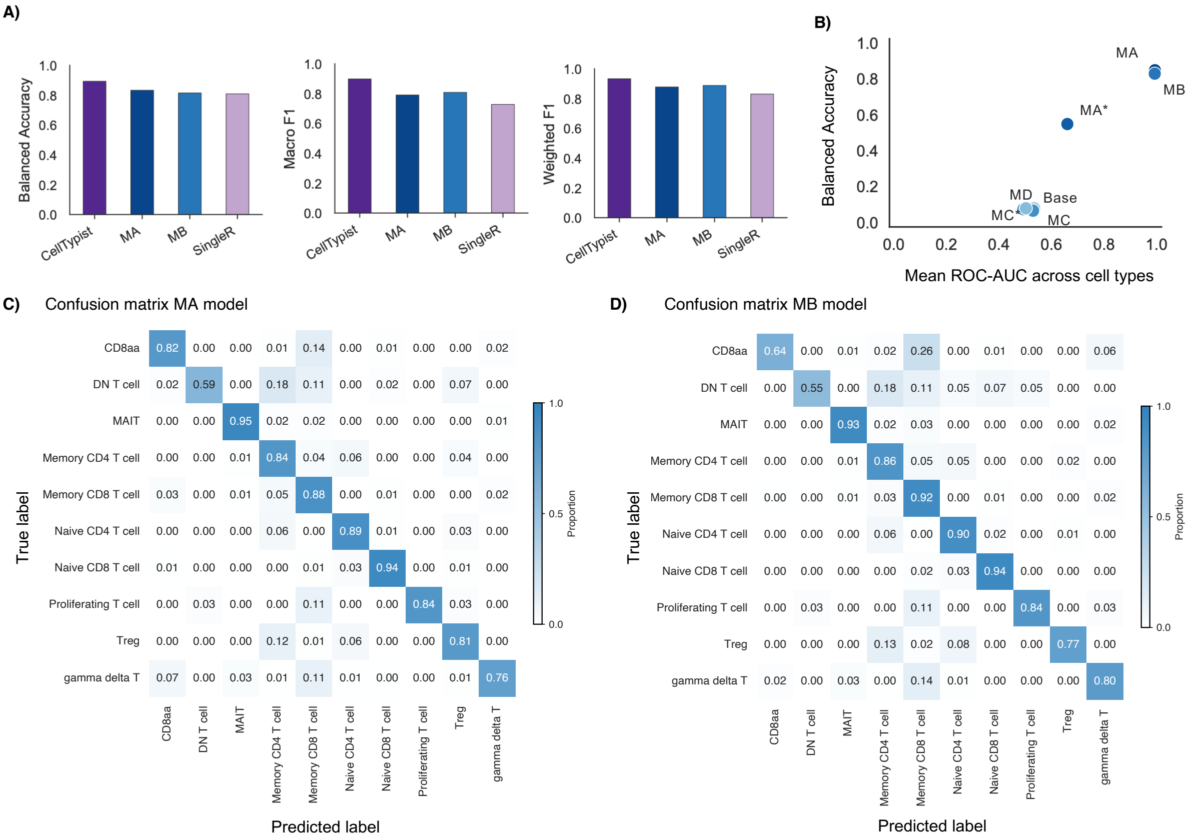


#### Embeddings

**E) Ablation study:**

| Method | Bio conservation: Isolated labels | KMeans NMI | KMeans ARI | Silhouette label | cLISI | Batch correction: BRAS | iLISI | KBET | Graph connectivity | PCR comparison | Aggregate score: Batch correction | Bio conservation | Total |
|---|---|---|---|---|---|---|---|---|---|---|---|---|---|
| **MB** | 0.63 | 0.72 | 0.75 | 0.81 | 0.99 | 0.72 | 0.27 | 0.90 | 0.77 | 0.51 | 0.63 | 0.78 | 0.72 |
| **MC** | 0.68 | 0.72 | 0.70 | 0.81 | 1.00 | 0.68 | 0.26 | 0.90 | 0.78 | 0.51 | 0.63 | 0.78 | 0.72 |
| **MA** | 0.62 | 0.68 | 0.71 | 0.75 | 0.99 | 0.78 | 0.27 | 0.92 | 0.80 | 0.56 | 0.66 | 0.75 | 0.72 |
| **MA*** | 0.57 | 0.65 | 0.68 | 0.75 | 0.99 | 0.79 | 0.27 | 0.91 | 0.83 | 0.53 | 0.66 | 0.73 | 0.70 |
| **Base** | 0.49 | 0.21 | 0.12 | 0.50 | 0.89 | 0.93 | 0.24 | 0.65 | 0.45 | 0.64 | 0.58 | 0.44 | 0.50 |
| **MD** | 0.49 | 0.05 | 0.03 | 0.45 | 0.81 | 0.87 | 0.27 | 0.63 | 0.29 | 0.85 | 0.58 | 0.37 | 0.45 |
| **MC*** | 0.46 | 0.02 | 0.01 | 0.46 | 0.76 | 0.91 | 0.30 | 0.37 | 0.12 | 0.99 | 0.54 | 0.34 | 0.42 |

**F) Benchmark:**

| Method | Bio conservation: Isolated labels | KMeans NMI | KMeans ARI | Silhouette label | cLISI | Batch correction: BRAS | iLISI | KBET | Graph connectivity | PCR comparison | Aggregate score: Batch correction | Bio conservation | Total |
|---|---|---|---|---|---|---|---|---|---|---|---|---|---|
| **MB** | 0.63 | 0.72 | 0.75 | 0.81 | 0.99 | 0.72 | 0.27 | 0.90 | 0.77 | 0.51 | 0.63 | 0.78 | 0.72 |
| **MA** | 0.62 | 0.68 | 0.71 | 0.75 | 0.99 | 0.78 | 0.27 | 0.92 | 0.80 | 0.56 | 0.66 | 0.75 | 0.72 |
| **PCA_onehot** | 0.52 | 0.43 | 0.26 | 0.51 | 0.94 | 0.94 | 0.22 | 0.69 | 0.72 | 0.54 | 0.62 | 0.53 | 0.57 |
| **PCA_rank** | 0.53 | 0.39 | 0.24 | 0.51 | 0.95 | 0.90 | 0.21 | 0.78 | 0.72 | 0.39 | 0.60 | 0.53 | 0.56 |
| **PCA_expression** | 0.53 | 0.50 | 0.35 | 0.51 | 0.99 | 0.87 | 0.17 | 0.56 | 0.79 | 0.00 | 0.48 | 0.58 | 0.54 |
| **scGPT** | 0.55 | 0.37 | 0.20 | 0.50 | 0.94 | 0.89 | 0.24 | 0.70 | 0.63 | 0.31 | 0.55 | 0.51 | 0.53 |
| **scVI_pretrain** | 0.56 | 0.32 | 0.19 | 0.51 | 0.95 | 0.80 | 0.22 | 0.72 | 0.72 | 0.00 | 0.49 | 0.51 | 0.50 |
| **scVI_adapted** | 0.56 | 0.32 | 0.19 | 0.51 | 0.95 | 0.80 | 0.22 | 0.72 | 0.72 | 0.00 | 0.49 | 0.51 | 0.50 |

**Figure S3: Cell-type annotation and embedding benchmark results for the *HIAI* T-cell dataset. A)** Cell type annotation benchmark results: Balanced accuracy, macro F1 score and weighted F1 score are reported for dataset-adapted CellTypist and SingleR and compared with model MA, trained on scRNA-seq-derived cell sentences, metadata labels, and literature-derived text, and model MB, trained on cell sentences and metadata labels without literature. **B)** Relationship between mean ROC-AUC across cell types (when evaluating performance for each cell type with a one-vs-rest-strategy) and balanced accuracy using all models included in the ablation study. **C), D)** Confusion matrices for cell-type annotation using the full model MA and the model without literature MB, respectively. **E), F)** Full single-cell integration benchmarking metric reports (following [2]) for the ablation models and benchmark/reference approaches, respectively.

#### Functionality annotation

**A) Functionality descriptions**

| Cell type | Category | Functionality description | Source |
|---|---|---|---|
| **Gamma delta T** | **True** | Recognize stress ligands independently of classical MHC molecules. | Vantourout & Hayday 2013 |
| | **True** | Recognize lipids, phosphoantigens, and stress ligands via non-conventional pathways. | |
| **Treg** | **True** | Produce immunosuppressive cytokines such as IL-10 and TGF-β | Vignali et al. 2008 |
| | **True** | Stably express FOXP3 to maintain suppressive identity. | |
| **MAIT** | **True** | Recognize bacterial-derived riboflavin metabolites presented on MR1. | Godfrey et al. 2019 |
| | **Paraphrase** | Detect bacterial riboflavin-derived metabolites displayed by MR1. | |
| | **Paraphrase** | Respond to microbial riboflavin metabolite antigens presented through MR1. | |
| | **Paraphrase** | Recognize MR1-presented metabolites generated by bacterial riboflavin biosynthesis. | |
| | **True** | Bridge innate and adaptive immunity with semi-invariant TCR and rapid cytokine output. | |
| | **Paraphrase** | Link innate and adaptive immune responses through a semi-invariant TCR and rapid cytokine production. | |
| | **Paraphrase** | Combine innate-like responsiveness with antigen-specific recognition via a semi-invariant TCR and fast cytokine release. | |
| | **Paraphrase** | Mediate rapid cytokine responses using a semi-invariant TCR at the interface of innate and adaptive immunity. | |
| **Memory CD8** | **True** | Rapidly kill target cells upon antigen re-exposure. | Weng et al. 2012 |
| | **True** | Retain long-term survival and recall potential | |
| **Memory CD4** | **True** | Effector memory subset lacks CCR7 and homes to inflamed tissues. | Weng et al. 2012 |
| | **True** | Central memory subset homes to lymphoid tissues and proliferates upon recall. | |
| **Naive CD4** | **True** | Serve as unprimed precursors to all helper T cell subsets. | Chapman et al. 2020 |
| | **True** | Act as a pool for generating novel responses to previously unseen antigens. | |
| **Proliferating T** | **True** | Undergo robust IL-2-driven expansion during acute response. | Chapman et al. 2020 |
| | **True** | Increased metabolic activity and cell cycle progression. | |

**B)** Cell type - functionality description: similarity scores

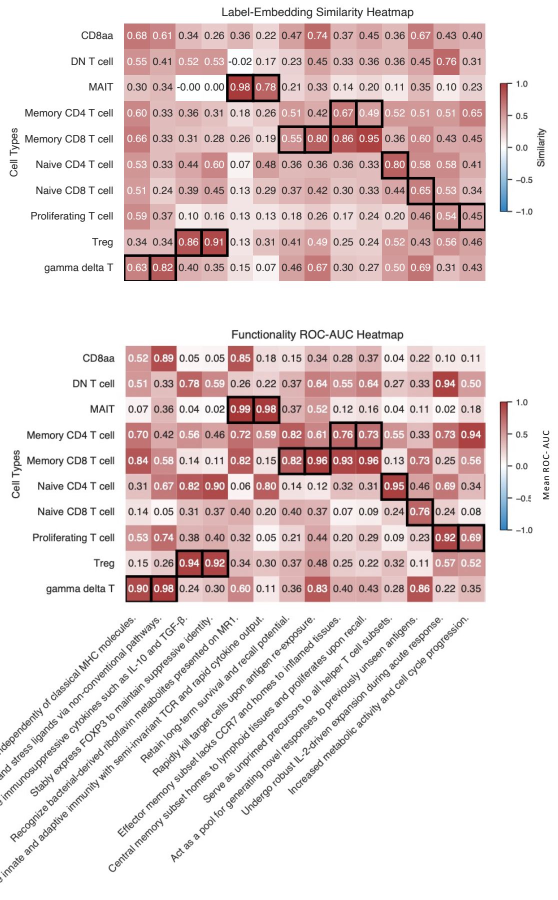


**Figure S4: Expert-curated T-cell functionality descriptions and their alignment in the full model. A)** Overview of all expert-curated functionality descriptions grouped by assigned T-cell type. Descriptions in the "true" category were used for the ablation and benchmark analyses, whereas "paraphrase" descriptions were to assess robustness to alternative wording in the ablation study results only. **B)** Heatmaps showing the cell-type label similarity scores and the cell-level mean ROC-AUC values for all functionality descriptions using the full model MA.

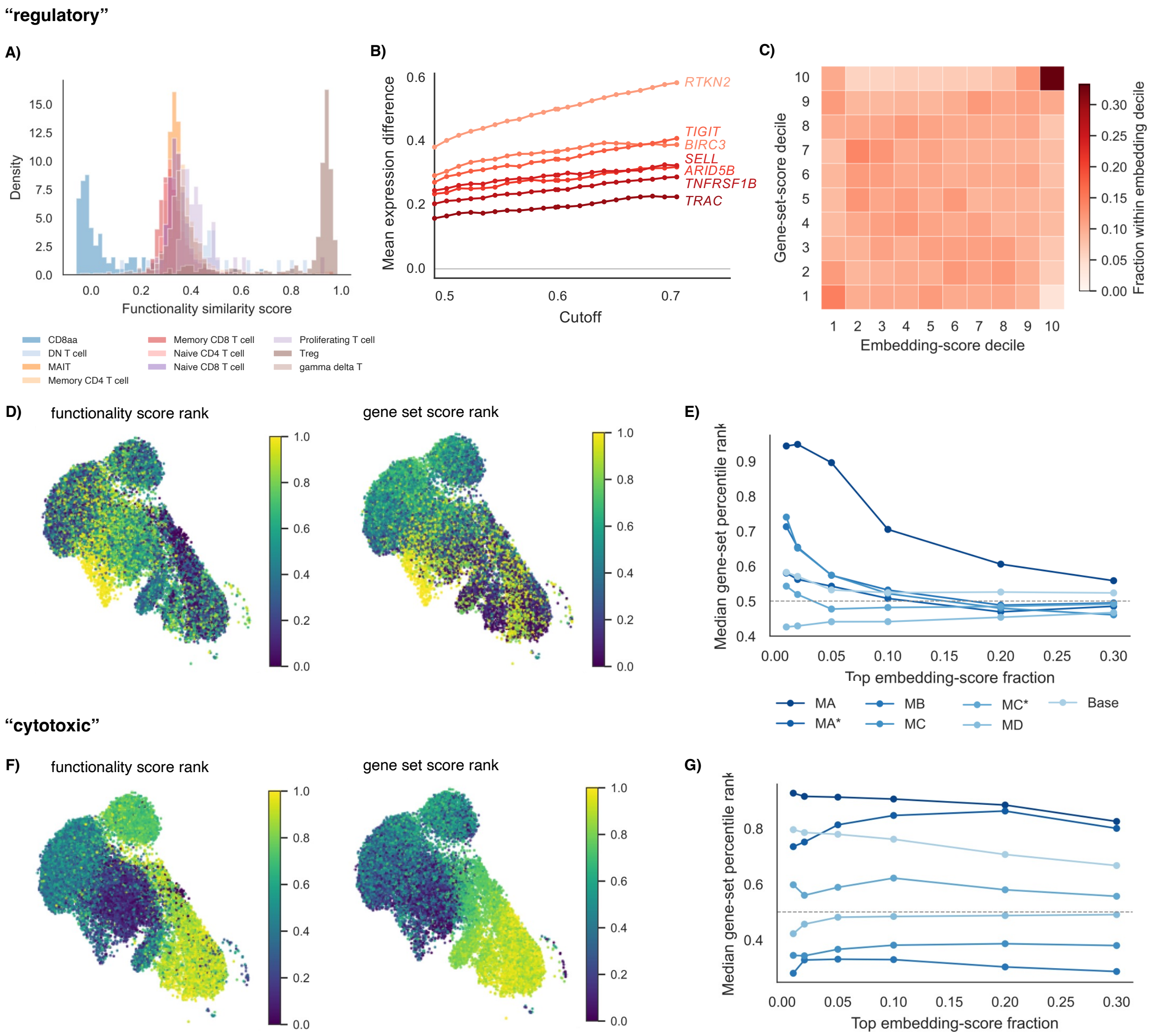


**Figure S5: Regulatory and cytotoxic functional query analyses.** **A)** Histogram of cell-wise similarity scores to the functionality annotation ‘regulatory’, colored by reference cell-type label, showing separation of cells with known regulatory identity. **B)** Sensitivity of differential expressed results of cells annotated as regulatory vs non-regulatory cells to the decision threshold, to assess robust identification of known markers. **C)** Concordance between embedding-based regulatory similarity scores and a regulatory gene set score from an established gene set after binning cells into score deciles. **D)** Regulatory similarity-score ranks and regulatory gene-set score ranks shown on the original gene expression based UMAP representation. **E)** Median regulatory gene-set percentile rank among cells in the top fractions of regulatory similarity scores, shown across ablation models. **F)** Cytotoxicity similarity-score ranks and cytotoxicity gene-set score ranks shown on the original gene expression based UMAP representation. **G)** Median cytotoxicity gene-set percentile rank among cells in the top fractions of cytotoxicity similarity scores, shown across ablation models.

**Disease interaction**

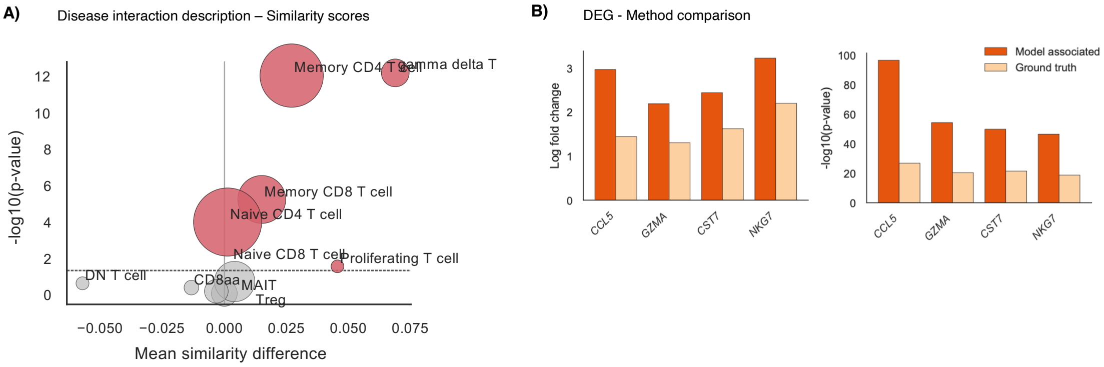


**Figure S6: CMV-associated similarity analysis without CMV-specific literature retrieval. A)** Separation of CMV status based on cell-wise cosine similarity to the disease-interaction description using the full MA model, trained without CMV metadata in the cell sentences and without dedicated CMV-specific literature retrieval. Cells were classified as CMV-associated when their similarity score exceeded the similarity threshold determined based on Youden's J (see Methods Section 5.14). Separation was evaluated within each cell type using the mean similarity difference and p-value from Mann-Whitney U test. Dot size indicates the number of cells per type. **B)** Comparison of log fold changes (logFC) and corresponding log-transformed exploratory $p$-values from differential gene expression (DEG) analysis of disease associated cells 'Memory CD4 T' cells separated either by similarity scores based on the model embeddings or by the CMV metadata labels.

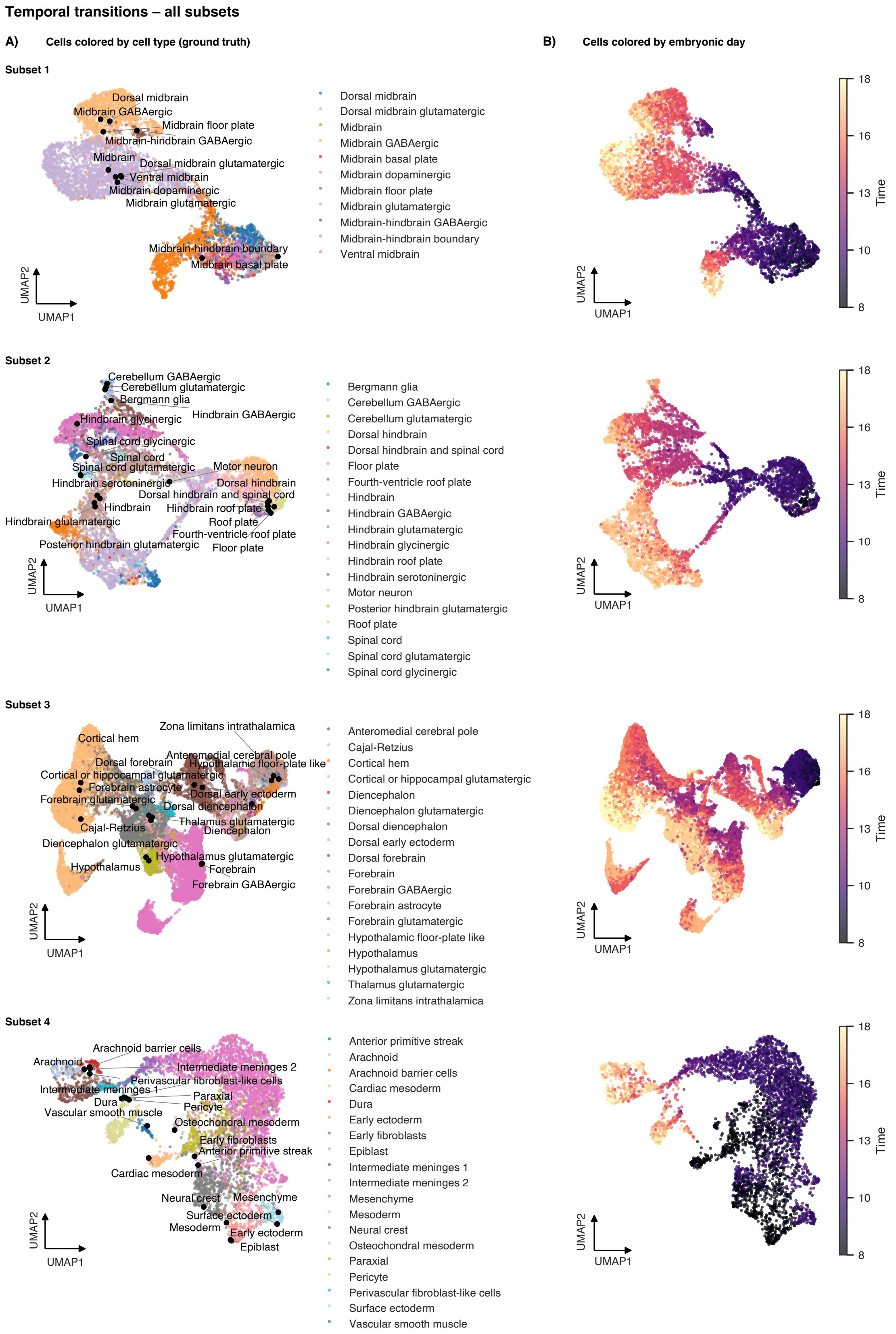


**Figure S7: Cell-type and temporal organization across all subsets of the LaManno embryonic mouse brain dataset (part 1). A)** Cells are colored by ground truth cell-type label with respective cell type labels in black. **B)** Cells are colored by temporal meta-data (embryonic day).

#### Temporal transitions – all subsets (continuation)

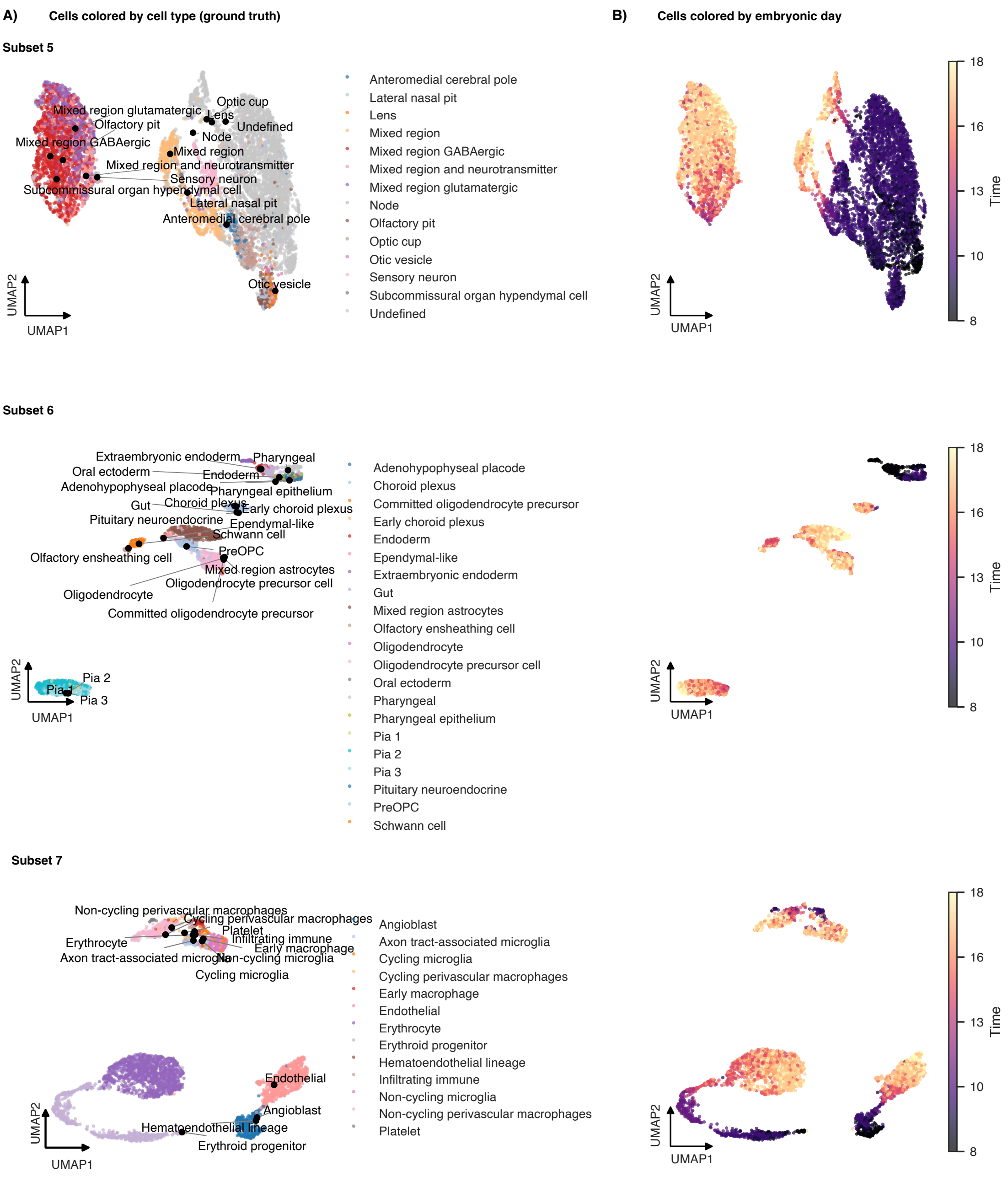


**Figure S8: Cell-type and temporal organization across all subsets of the LaManno embryonic mouse brain dataset (continued). A)** Cells are colored by ground truth cell-type label with respective cell type labels in black. **B)** Cells are colored by temporal meta-data (embryonic day).

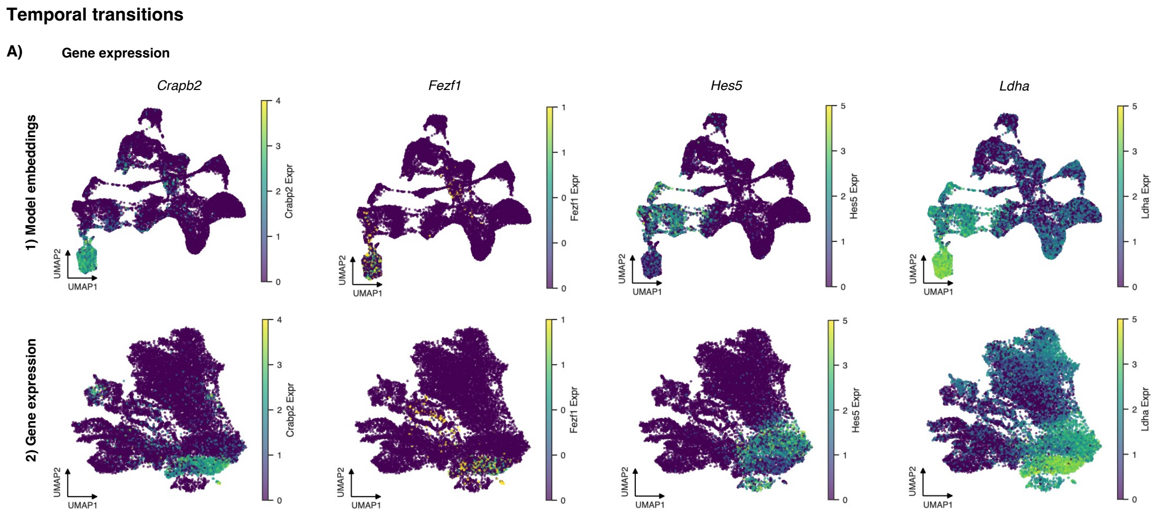


**Figure S9: Marker gene expression along the forebrain developmental trajectory.** Expression of *Crabp2*, *Dlk1*, *Fezf1*, and *Hes5* is shown on UMAP representations based on model embeddings and on the original gene expression data. The displayed cells correspond to the forebrain trajectory analyzed in Fig. 5, from ‘Dorsal forebrain’ to ‘Cortical or hippocampal glutamatergic’ and ‘Forebrain GABAergic’ cells.

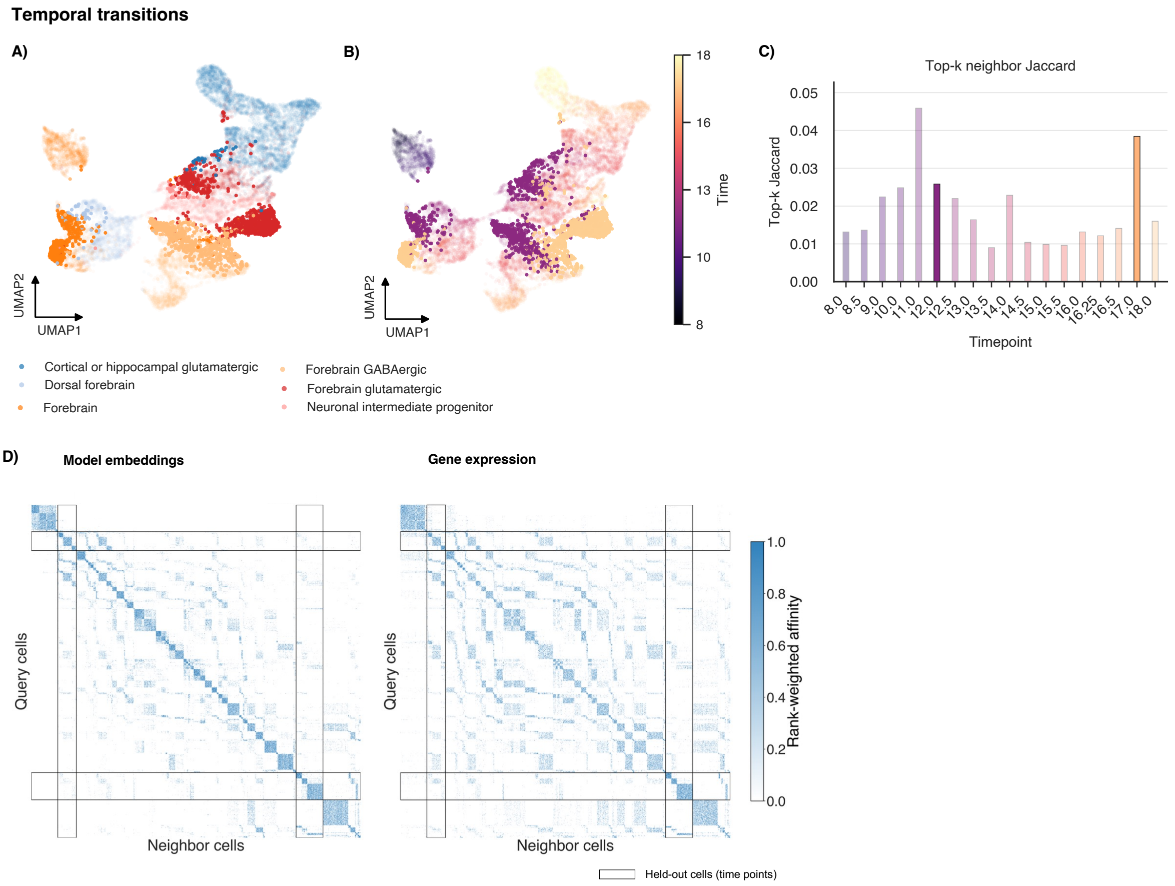


**Figure S10: Projection of held-out embryonic time points. A), B)**: UMAP representations of the forebrain trajectory using a model trained with selected embryonic time points withheld. Cells are colored by reference cell type in **A)** and by embryonic day in **B)**, with cells from held-out time points highlighted. **C)** Top-$k$ neighbor Jaccard similarity comparing neighborhoods in model embeddings and expression-based embeddings across time points. **D)** Rank-weighted affinity of single cells in model embeddings and expression-based embeddings, with held-out cells indicated by black rectangles.
