## Additional file 2 for "Adding layers of information to scRNA-seq data using pre-trained language models"

### Additional file 2 for
### Adding layers of information to scRNA-seq data using pre-trained language models

Sonia Maria Krißmer ✉[1,2], Jonatan Menger[1,2], Johan Rollin[3], Tanja Vogel[3], Harald Binder[1,2,4], Maren Hackenberg ✉[1,4,5]

[1] Institute of Medical Biometry and Statistics (IMBI), Faculty of Medicine and Medical Center, University of Freiburg, Germany
[2] CIBSS – Centre for Integrative Biological Signalling Studies, University of Freiburg, Germany
[3] Institute of Anatomy and Cell Biology, Department Molecular Embryology, Faculty of Medicine, University of Freiburg, Germany
[4] Freiburg Center for Data Analysis, Modeling and AI, University of Freiburg, Germany
[5] Present address: European Molecular Biology Laboratory, Heidelberg, Germany

August 10, 2026

### Supplementary reports

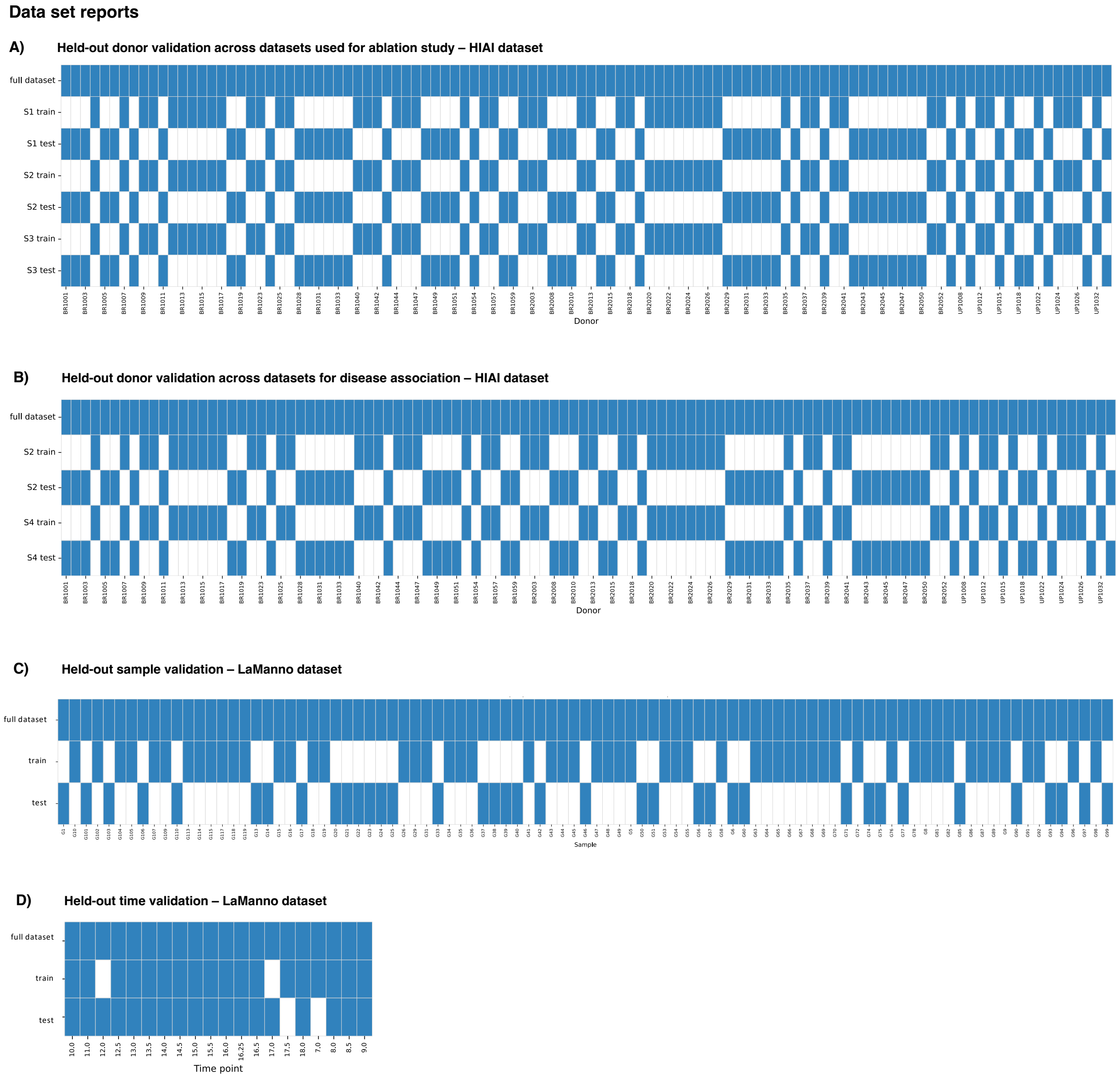


**Figure R1: Training and evaluation splits used across datasets. A)** Held-out donor splits for the HIAI datasets S1, S2 and S3 required for the training of the ablation study models. **B)** Held-out donor splits for disease interaction results (HIAI dataset S4 including CMV metadata) using the HIAI dataset. For comparison with results using the MA model, S2 donor split is again shown here. **C)** Held-out mouse split used for the developmental trajectory analysis on the LaManno dataset. **D)** Held-out embryonic time point split used for the supplementary temporal generalization analysis on the LaManno dataset.

**Table R1:** Dataset report for datasets required for training all models used as part of the ablation study. Report includes cell type proportion per cell type in the full dataset as well as train and test split for scRNA-seq-based datasets and literature search results per cell type for literature-based datasets.

<table>
<tr><th>Dataset ID</th><th>Modality</th><th>Split / construction</th><th>Semantic labels</th><th>Shuffled labels</th><th>Report</th><th>Models</th></tr>
<tr><td>S1</td><td>scRNA-seq</td><td>Held-out donor split</td><td>Yes</td><td>No</td><td>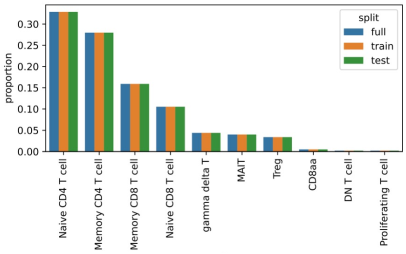
</td><td>MA<br>MA*<br>MB</td></tr>
<tr><td>S2</td><td>scRNA-seq</td><td>Held-out donor split with nonsemantic cell sentences</td><td>No</td><td>No</td><td>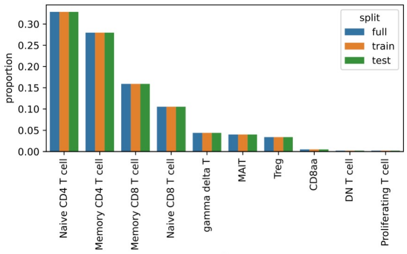
</td><td>MC</td></tr>
<tr><td>S3</td><td>scRNA-seq</td><td>Held-out donor split with shuffled cell-type labels</td><td>No</td><td>Yes</td><td>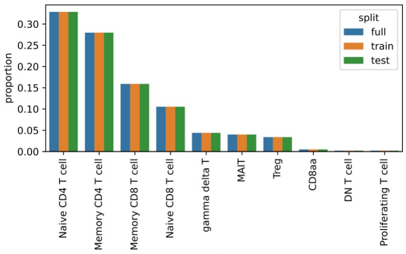
</td><td>MC*</td></tr>
<tr><td>N1</td><td>literature</td><td>Literature corpus</td><td>NA</td><td>No</td><td>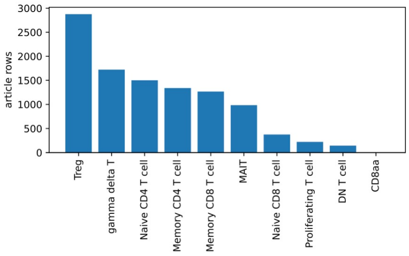
</td><td>MA<br>MD</td></tr>
<tr><td>N2</td><td>literature</td><td>Literature corpus with shuffled labels</td><td>NA</td><td>Yes</td><td>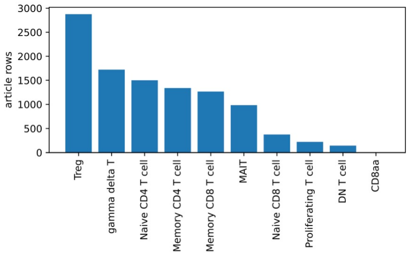
</td><td>MA*</td></tr>
</table>

**Table R2 :** PubMed PMID and CL term retrieval and inclusion summary for N1 and N2 NCBI literature datasets for the ablation study based on the *HIAI* dataset.

| Query label | Search query | PubMed count | Returned | Retrieved | N1/N2 included | CL/scType rows | CL/scType training rows |
|---|---|---|---|---|---|---|---|
| CD8aa | (homo sapiens[Mesh]) AND (Blood Cells[Mesh]) AND ("CD8aa"[Title/Abstract] OR "CD8aas"[Title/Abstract]) | 2 | 2 | 2 | 1 | 18 | 36 |
| DN T cell | (homo sapiens[Mesh]) AND (Blood Cells[Mesh]) AND ("DN T cell"[Title/Abstract] OR "DN T cells"[Title/Abstract]) | 160 | 160 | 160 | 144 | 2 | 4 |
| MAIT | (homo sapiens[Mesh]) AND (Blood Cells[Mesh]) AND ("MAIT"[Title/Abstract] OR "MAITs"[Title/Abstract]) | 1,029 | 1,029 | 1,029 | 984 | 2 | 4 |
| Memory CD4 T cell | (homo sapiens[Mesh]) AND (Blood Cells[Mesh]) AND ("Memory CD4 T cell"[Title/Abstract] OR "Memory CD4 T cells"[Title/Abstract]) | 1,660 | 1,660 | 1,660 | 1,340 | 2 | 4 |
| Memory CD8 T cell | (homo sapiens[Mesh]) AND (Blood Cells[Mesh]) AND ("Memory CD8 T cell"[Title/Abstract] OR "Memory CD8 T cells"[Title/Abstract]) | 1,483 | 1,483 | 1,483 | 1,267 | 2 | 4 |
| Naive CD4 T cell | (homo sapiens[Mesh]) AND (Blood Cells[Mesh]) AND ("Naive CD4 T cell"[Title/Abstract] OR "Naive CD4 T cells"[Title/Abstract]) | 1,843 | 1,843 | 1,843 | 1,501 | 2 | 4 |
| Naive CD8 T cell | (homo sapiens[Mesh]) AND (Blood Cells[Mesh]) AND ("Naive CD8 T cell"[Title/Abstract] OR "Naive CD8 T cells"[Title/Abstract]) | 558 | 558 | 558 | 375 | 2 | 4 |
| Proliferating T cell | (homo sapiens[Mesh]) AND (Blood Cells[Mesh]) AND ("Proliferating T cell"[Title/Abstract] OR "Proliferating T cells"[Title/Abstract]) | 230 | 230 | 230 | 222 | 2 | 4 |
| Treg | (homo sapiens[Mesh]) AND (Blood Cells[Mesh]) AND ("Treg"[Title/Abstract] OR "Tregs"[Title/Abstract]) | 18,252 | 3,000 | 3,000 | 2,876 | 15 | 30 |
| gamma delta T | (homo sapiens[Mesh]) AND (Blood Cells[Mesh]) AND ("gamma delta T"[Title/Abstract] OR "gamma delta Ts"[Title/Abstract]) | 1,773 | 1,773 | 1,773 | 1,722 | 15 | 30 |

**Table R3 :** PubMed PMID retrieval and inclusion summary for the N3 literature-based dataset for the HIAI T cell dataset tailored to CMV disease interactions.

| Query label | Search query | PubMed count | Returned | Retrieved | N4 included | CL/scType rows | CL/scType training rows |
|---|---|---|---|---|---|---|---|
| CD8aa | ((homo sapiens[Mesh]) AND (Blood Cells[Mesh]) AND ("CD8aa"[Title/Abstract] OR "CD8aas"[Title/Abstract]) NOT CMV) | 2 | 2 | 2 | 1 | 18 | 36 |
| DN T cell | ((homo sapiens[Mesh]) AND (Blood Cells[Mesh]) AND ("DN T cell"[Title/Abstract] OR "DN T cells"[Title/Abstract]) NOT CMV) | 157 | 157 | 157 | 141 | 2 | 4 |
| DN T cell_CMV | ((homo sapiens[Mesh]) AND (Blood Cells[Mesh]) AND ("DN T cell"[Title/Abstract] OR "DN T cells"[Title/Abstract]) AND CMV) | 3 | 3 | 3 | 3 | 2 | 4 |
| MAIT | ((homo sapiens[Mesh]) AND (Blood Cells[Mesh]) AND ("MAIT"[Title/Abstract] OR "MAITs"[Title/Abstract]) NOT CMV) | 1,025 | 1,025 | 1,025 | 980 | 2 | 4 |
| MAIT_CMV | ((homo sapiens[Mesh]) AND (Blood Cells[Mesh]) AND ("MAIT"[Title/Abstract] OR "MAITs"[Title/Abstract]) AND CMV) | 6 | 6 | 6 | 5 | 2 | 4 |
| Memory CD4 T cell | ((homo sapiens[Mesh]) AND (Blood Cells[Mesh]) AND ("Memory CD4 T cell"[Title/Abstract] OR "Memory CD4 T cells"[Title/Abstract]) NOT CMV) | 1,622 | 1,622 | 1,622 | 1,305 | 2 | 4 |
| Memory CD4 T cell_CMV | ((homo sapiens[Mesh]) AND (Blood Cells[Mesh]) AND ("Memory CD4 T cell"[Title/Abstract] OR "Memory CD4 T cells"[Title/Abstract]) AND CMV) | 40 | 40 | 40 | 34 | 2 | 4 |
| Memory CD8 T cell | ((homo sapiens[Mesh]) AND (Blood Cells[Mesh]) AND ("Memory CD8 T cell"[Title/Abstract] OR "Memory CD8 T cells"[Title/Abstract]) NOT CMV) | 1,417 | 1,417 | 1,417 | 1,209 | 2 | 4 |
| Memory CD8 T cell_CMV | ((homo sapiens[Mesh]) AND (Blood Cells[Mesh]) AND ("Memory CD8 T cell"[Title/Abstract] OR "Memory CD8 T cells"[Title/Abstract]) AND CMV) | 70 | 70 | 70 | 62 | 2 | 4 |
| Naive CD4 T cell | ((homo sapiens[Mesh]) AND (Blood Cells[Mesh]) AND ("Naive CD4 T cell"[Title/Abstract] OR "Naive CD4 T cells"[Title/Abstract]) NOT CMV) | 1,830 | 1,830 | 1,830 | 1,492 | 2 | 4 |
| Naive CD4 T cell_CMV | ((homo sapiens[Mesh]) AND (Blood Cells[Mesh]) AND ("Naive CD4 T cell"[Title/Abstract] OR "Naive CD4 T cells"[Title/Abstract]) AND CMV) | 16 | 16 | 16 | 9 | 2 | 4 |
| Naive CD8 T cell | ((homo sapiens[Mesh]) AND (Blood Cells[Mesh]) AND ("Naive CD8 T cell"[Title/Abstract] OR "Naive CD8 T cells"[Title/Abstract]) NOT CMV) | 528 | 528 | 528 | 355 | 2 | 4 |
| Naive CD8 T cell_CMV | ((homo sapiens[Mesh]) AND (Blood Cells[Mesh]) AND ("Naive CD8 T cell"[Title/Abstract] OR "Naive CD8 T cells"[Title/Abstract]) AND CMV) | 30 | 30 | 30 | 20 | 2 | 4 |
| Proliferating T cell | ((homo sapiens[Mesh]) AND (Blood Cells[Mesh]) AND ("Proliferating T cell"[Title/Abstract] OR "Proliferating T cells"[Title/Abstract]) NOT CMV) | 227 | 227 | 227 | 219 | 2 | 4 |
| Proliferating T cell_CMV | ((homo sapiens[Mesh]) AND (Blood Cells[Mesh]) AND ("Proliferating T cell"[Title/Abstract] OR "Proliferating T cells"[Title/Abstract]) AND CMV) | 3 | 3 | 3 | 2 | 2 | 4 |

12 Continued on next page

**Table R3 :** PubMed PMID retrieval and inclusion summary for the N3 literature-based dataset for the HIAI T cell dataset tailored to CMV disease interactions. (Continued)

| Query label | Search query | PubMed count | Returned | Retrieved | N4 included | CL/scType rows | CL/scType training rows |
|---|---|---|---|---|---|---|---|
| Treg | ((homo sapiens[Mesh]) AND (Blood Cells[Mesh]) AND ("Treg"[Title/Abstract] OR "Tregs"[Title/Abstract]) NOT CMV) | 18,208 | 3,000 | 3,000 | 2,877 | 15 | 30 |
| Treg_CMV | ((homo sapiens[Mesh]) AND (Blood Cells[Mesh]) AND ("Treg"[Title/Abstract] OR "Tregs"[Title/Abstract]) AND CMV) | 78 | 78 | 78 | 70 | 15 | 30 |
| gamma delta T | ((homo sapiens[Mesh]) AND (Blood Cells[Mesh]) AND ("gamma delta T"[Title/Abstract] OR "gamma delta Ts"[Title/Abstract]) NOT CMV) | 1,759 | 1,759 | 1,759 | 1,709 | 15 | 30 |
| gamma delta T_CMV | ((homo sapiens[Mesh]) AND (Blood Cells[Mesh]) AND ("gamma delta T"[Title/Abstract] OR "gamma delta Ts"[Title/Abstract]) AND CMV) | 14 | 14 | 14 | 13 | 15 | 30 |

**Table R4 :** PubMed PMID retrieval and inclusion summary for the N4 NCBI literature datasets for the *LaManno* dataset.

| Query label | Search query | PubMed count | Returned | Retrieved | N4 included |
|---|---|---|---|---|---|
| Adenohypophyseal placode | (mouse[Mesh]) AND (Embryo, Mammalian[Mesh]) AND ("Adenohypophyseal placode"[Title/Abstract] OR "Adenohypophyseal placodes"[Title/Abstract]) | 1 | 1 | 1 | 0 |
| Angioblast | (mouse[Mesh]) AND (Embryo, Mammalian[Mesh]) AND ("Angioblast"[Title/Abstract] OR "Angioblasts"[Title/Abstract]) | 30 | 30 | 30 | 3 |
| Anterior | (mouse[Mesh]) AND (Embryo, Mammalian[Mesh]) AND ("Anterior"[Title/Abstract] OR "Anteriors"[Title/Abstract]) | 658 | 658 | 648 | 208 |
| Anterior primitive streak | (mouse[Mesh]) AND (Embryo, Mammalian[Mesh]) AND ("Anterior primitive streak"[Title/Abstract] OR "Anterior primitive streaks"[Title/Abstract]) | 10 | 10 | 10 | 0 |
| Arachnoid | (mouse[Mesh]) AND (Embryo, Mammalian[Mesh]) AND ("Arachnoid"[Title/Abstract] OR "Arachnoids"[Title/Abstract]) | 1 | 1 | 1 | 0 |
| Bergmann glia | (mouse[Mesh]) AND (Embryo, Mammalian[Mesh]) AND ("Bergmann glia"[Title/Abstract] OR "Bergmann glias"[Title/Abstract]) | 13 | 13 | 13 | 7 |
| Cajal-Retzius | (mouse[Mesh]) AND (Embryo, Mammalian[Mesh]) AND ("Cajal-Retzius"[Title/Abstract]) | 39 | 39 | 39 | 23 |
| Cardiac | (mouse[Mesh]) AND (Embryo, Mammalian[Mesh]) AND ("Cardiac"[Title/Abstract] OR "Cardiacs"[Title/Abstract]) | 834 | 834 | 824 | 505 |
| Cardiac mesoderm | (mouse[Mesh]) AND (Embryo, Mammalian[Mesh]) AND ("Cardiac mesoderm"[Title/Abstract] OR "Cardiac mesoderms"[Title/Abstract]) | 22 | 22 | 22 | 0 |
| Caudal | (mouse[Mesh]) AND (Embryo, Mammalian[Mesh]) AND ("Caudal"[Title/Abstract] OR "Caudals"[Title/Abstract]) | 284 | 284 | 284 | 103 |
| Chondrocytes | (mouse[Mesh]) AND (Embryo, Mammalian[Mesh]) AND ("Chondrocytes"[Title/Abstract]) | 214 | 214 | 214 | 172 |
| Choroid plexus | (mouse[Mesh]) AND (Embryo, Mammalian[Mesh]) AND ("Choroid plexus"[Title/Abstract]) | 72 | 72 | 72 | 31 |
| Cortical hem | (mouse[Mesh]) AND (Embryo, Mammalian[Mesh]) AND ("Cortical hem"[Title/Abstract] OR "Cortical hems"[Title/Abstract]) | 18 | 18 | 18 | 1 |

Continued on next page

**Table R4 :** PubMed PMID retrieval and inclusion summary for the N4 NCBI literature datasets for the *LaManno* dataset. (Continued)

| Query label | Search query | PubMed count | Returned | Retrieved | N4 included |
|---|---|---|---|---|---|
| Definitive endoderm | (mouse[Mesh]) AND (Embryo, Mammalian[Mesh]) AND ("Definitive endoderm"[Title/Abstract] OR "Definitive endoderms"[Title/Abstract]) | 73 | 73 | 73 | 0 |
| Diencephalic roof plate | (mouse[Mesh]) AND (Embryo, Mammalian[Mesh]) AND ("Diencephalic roof plate"[Title/Abstract] OR "Diencephalic roof plates"[Title/Abstract]) | 1 | 1 | 1 | 0 |
| Diencephalon | (mouse[Mesh]) AND (Embryo, Mammalian[Mesh]) AND ("Diencephalon"[Title/Abstract] OR "Diencephalons"[Title/Abstract]) | 75 | 75 | 75 | 13 |
| Dorsal diencephalon | (mouse[Mesh]) AND (Embryo, Mammalian[Mesh]) AND ("Dorsal diencephalon"[Title/Abstract] OR "Dorsal diencephalons"[Title/Abstract]) | 1 | 1 | 1 | 0 |
| Dorsal forebrain | (mouse[Mesh]) AND (Embryo, Mammalian[Mesh]) AND ("Dorsal forebrain"[Title/Abstract] OR "Dorsal forebrains"[Title/Abstract]) | 6 | 6 | 6 | 1 |
| Dorsal hindbrain | (mouse[Mesh]) AND (Embryo, Mammalian[Mesh]) AND ("Dorsal hindbrain"[Title/Abstract] OR "Dorsal hindbrains"[Title/Abstract]) | 2 | 2 | 2 | 0 |
| Dorsal midbrain | (mouse[Mesh]) AND (Embryo, Mammalian[Mesh]) AND ("Dorsal midbrain"[Title/Abstract] OR "Dorsal midbrains"[Title/Abstract]) | 5 | 5 | 5 | 0 |
| Early ectoderm | (mouse[Mesh]) AND (Embryo, Mammalian[Mesh]) AND ("Early ectoderm"[Title/Abstract] OR "Early ectoderms"[Title/Abstract]) | 2 | 2 | 2 | 0 |
| Early fibroblasts | (mouse[Mesh]) AND (Embryo, Mammalian[Mesh]) AND ("Early fibroblasts"[Title/Abstract]) | 3 | 3 | 3 | 3 |
| Early macrophage | (mouse[Mesh]) AND (Embryo, Mammalian[Mesh]) AND ("Early macrophage"[Title/Abstract] OR "Early macrophages"[Title/Abstract]) | 2 | 2 | 2 | 2 |
| Endoderm | (mouse[Mesh]) AND (Embryo, Mammalian[Mesh]) AND ("Endoderm"[Title/Abstract] OR "Endoderms"[Title/Abstract]) | 1,078 | 1,078 | 1,068 | 198 |
| Endothelial | (mouse[Mesh]) AND (Embryo, Mammalian[Mesh]) AND ("Endothelial"[Title/Abstract] OR "Endothelials"[Title/Abstract]) | 1,453 | 1,453 | 1,430 | 959 |
| Epiblast | (mouse[Mesh]) AND (Embryo, Mammalian[Mesh]) AND ("Epiblast"[Title/Abstract] OR "Epiblasts"[Title/Abstract]) | 697 | 697 | 697 | 282 |
| Erythrocyte | (mouse[Mesh]) AND (Embryo, Mammalian[Mesh]) AND ("Erythrocyte"[Title/Abstract] OR "Erythrocytes"[Title/Abstract]) | 167 | 167 | 167 | 117 |
| Erythroid progenitor | (mouse[Mesh]) AND (Embryo, Mammalian[Mesh]) AND ("Erythroid progenitor"[Title/Abstract] OR "Erythroid progenitors"[Title/Abstract]) | 62 | 62 | 62 | 37 |
| Extraembryonic ectoderm | (mouse[Mesh]) AND (Embryo, Mammalian[Mesh]) AND ("Extraembryonic ectoderm"[Title/Abstract] OR "Extraembryonic ectoderms"[Title/Abstract]) | 97 | 97 | 97 | 31 |
| Extraembryonic endoderm | (mouse[Mesh]) AND (Embryo, Mammalian[Mesh]) AND ("Extraembryonic endoderm"[Title/Abstract] OR "Extraembryonic endoderms"[Title/Abstract]) | 92 | 92 | 92 | 1 |
| Floor plate | (mouse[Mesh]) AND (Embryo, Mammalian[Mesh]) AND ("Floor plate"[Title/Abstract] OR "Floor plates"[Title/Abstract]) | 82 | 82 | 82 | 16 |
| Forebrain | (mouse[Mesh]) AND (Embryo, Mammalian[Mesh]) AND ("Forebrain"[Title/Abstract] OR "Forebrains"[Title/Abstract]) | 425 | 425 | 365 | 188 |
| Forebrain GABAergic | (mouse[Mesh]) AND (Embryo, Mammalian[Mesh]) AND ("Forebrain GABAergic"[Title/Abstract] OR "Forebrain GABAergics"[Title/Abstract]) | 1 | 1 | 1 | 0 |

16 Continued on next page

**Table R4 :** PubMed PMID retrieval and inclusion summary for the N4 NCBI literature datasets for the *LaManno* dataset. (Continued)

| Query label | Search query | PubMed count | Returned | Retrieved | N4 included |
|---|---|---|---|---|---|
| Forebrain astrocyte | (mouse[Mesh]) AND (Embryo, Mammalian[Mesh]) AND ("Forebrain astrocyte"[Title/Abstract] OR "Forebrain astrocytes"[Title/Abstract]) | 3 | 3 | 3 | 0 |
| Forebrain glutamatergic | (mouse[Mesh]) AND (Embryo, Mammalian[Mesh]) AND ("Forebrain glutamatergic"[Title/Abstract] OR "Forebrain glutamatergics"[Title/Abstract]) | 1 | 1 | 1 | 0 |
| Gut | (mouse[Mesh]) AND (Embryo, Mammalian[Mesh]) AND ("Gut"[Title/Abstract] OR "Guts"[Title/Abstract]) | 278 | 278 | 278 | 95 |
| Hindbrain | (mouse[Mesh]) AND (Embryo, Mammalian[Mesh]) AND ("Hindbrain"[Title/Abstract] OR "Hindbrains"[Title/Abstract]) | 240 | 240 | 180 | 37 |
| Hypothalamus | (mouse[Mesh]) AND (Embryo, Mammalian[Mesh]) AND ("Hypothalamus"[Title/Abstract]) | 108 | 108 | 108 | 58 |
| Infiltrating immune | (mouse[Mesh]) AND (Embryo, Mammalian[Mesh]) AND ("Infiltrating immune"[Title/Abstract] OR "Infiltrating immunes"[Title/Abstract]) | 1 | 1 | 1 | 1 |
| Lens | (mouse[Mesh]) AND (Embryo, Mammalian[Mesh]) AND ("Lens"[Title/Abstract]) | 137 | 137 | 137 | 75 |
| Mesenchyme | (mouse[Mesh]) AND (Embryo, Mammalian[Mesh]) AND ("Mesenchyme"[Title/Abstract] OR "Mesenchymes"[Title/Abstract]) | 589 | 589 | 469 | 223 |
| Mesoderm | (mouse[Mesh]) AND (Embryo, Mammalian[Mesh]) AND ("Mesoderm"[Title/Abstract] OR "Mesoderms"[Title/Abstract]) | 827 | 827 | 827 | 195 |
| Midbrain | (mouse[Mesh]) AND (Embryo, Mammalian[Mesh]) AND ("Midbrain"[Title/Abstract] OR "Midbrains"[Title/Abstract]) | 256 | 256 | 256 | 68 |
| Midbrain GABAergic | (mouse[Mesh]) AND (Embryo, Mammalian[Mesh]) AND ("Midbrain GABAergic"[Title/Abstract] OR "Midbrain GABAergics"[Title/Abstract]) | 1 | 1 | 1 | 0 |
| Midbrain dopaminergic | (mouse[Mesh]) AND (Embryo, Mammalian[Mesh]) AND ("Midbrain dopaminergic"[Title/Abstract] OR "Midbrain dopaminergics"[Title/Abstract]) | 44 | 44 | 44 | 0 |
| Midbrain floor plate | (mouse[Mesh]) AND (Embryo, Mammalian[Mesh]) AND ("Midbrain floor plate"[Title/Abstract] OR "Midbrain floor plates"[Title/Abstract]) | 2 | 2 | 2 | 0 |
| Midbrain-hindbrain boundary | (mouse[Mesh]) AND (Embryo, Mammalian[Mesh]) AND ("Midbrain-hindbrain boundary"[Title/Abstract] OR "Midbrain-hindbrain boundarys"[Title/Abstract]) | 16 | 16 | 16 | 0 |
| Motor neuron | (mouse[Mesh]) AND (Embryo, Mammalian[Mesh]) AND ("Motor neuron"[Title/Abstract] OR "Motor neurons"[Title/Abstract]) | 212 | 212 | 212 | 89 |
| Nascent mesoderm | (mouse[Mesh]) AND (Embryo, Mammalian[Mesh]) AND ("Nascent mesoderm"[Title/Abstract] OR "Nascent mesoderms"[Title/Abstract]) | 15 | 15 | 15 | 0 |
| Neural crest | (mouse[Mesh]) AND (Embryo, Mammalian[Mesh]) AND ("Neural crest"[Title/Abstract] OR "Neural crests"[Title/Abstract]) | 497 | 497 | 497 | 192 |
| Neuromesodermal progenitors | (mouse[Mesh]) AND (Embryo, Mammalian[Mesh]) AND ("Neuromesodermal progenitors"[Title/Abstract]) | 13 | 13 | 13 | 3 |
| Neuronal intermediate progenitor | (mouse[Mesh]) AND (Embryo, Mammalian[Mesh]) AND ("Neuronal intermediate progenitor"[Title/Abstract] OR "Neuronal intermediate progenitors"[Title/Abstract]) | 1 | 1 | 1 | 1 |
| Node | (mouse[Mesh]) AND (Embryo, Mammalian[Mesh]) AND ("Node"[Title/Abstract] OR "Nodes"[Title/Abstract]) | 247 | 247 | 247 | 136 |
| Olfactory ensheathing cell | (mouse[Mesh]) AND (Embryo, Mammalian[Mesh]) AND ("Olfactory ensheathing cell"[Title/Abstract] OR "Olfactory ensheathing cells"[Title/Abstract]) | 9 | 9 | 9 | 0 |

17 Continued on next page

**Table R4 :** PubMed PMID retrieval and inclusion summary for the N4 NCBI literature datasets for the *LaManno* dataset. (Continued)

| Query label | Search query | PubMed count | Returned | Retrieved | N4 included |
|---|---|---|---|---|---|
| Olfactory epithelium | (mouse[Mesh]) AND (Embryo, Mammalian[Mesh]) AND ("Olfactory epithelium"[Title/Abstract] OR "Olfactory epitheliums"[Title/Abstract]) | 78 | 78 | 78 | 18 |
| Olfactory pit | (mouse[Mesh]) AND (Embryo, Mammalian[Mesh]) AND ("Olfactory pit"[Title/Abstract] OR "Olfactory pits"[Title/Abstract]) | 7 | 7 | 7 | 2 |
| Oligodendrocyte | (mouse[Mesh]) AND (Embryo, Mammalian[Mesh]) AND ("Oligodendrocyte"[Title/Abstract] OR "Oligodendrocytes"[Title/Abstract]) | 214 | 214 | 214 | 119 |
| Oligodendrocyte precursor cell | (mouse[Mesh]) AND (Embryo, Mammalian[Mesh]) AND ("Oligodendrocyte precursor cell"[Title/Abstract] OR "Oligodendrocyte precursor cells"[Title/Abstract]) | 21 | 21 | 21 | 0 |
| Optic cup | (mouse[Mesh]) AND (Embryo, Mammalian[Mesh]) AND ("Optic cup"[Title/Abstract] OR "Optic cups"[Title/Abstract]) | 29 | 29 | 29 | 11 |
| Oral ectoderm | (mouse[Mesh]) AND (Embryo, Mammalian[Mesh]) AND ("Oral ectoderm"[Title/Abstract] OR "Oral ectoderms"[Title/Abstract]) | 8 | 8 | 8 | 2 |
| Otic vesicle | (mouse[Mesh]) AND (Embryo, Mammalian[Mesh]) AND ("Otic vesicle"[Title/Abstract] OR "Otic vesicles"[Title/Abstract]) | 44 | 44 | 44 | 16 |
| Paraxial | (mouse[Mesh]) AND (Embryo, Mammalian[Mesh]) AND ("Paraxial"[Title/Abstract] OR "Paraxials"[Title/Abstract]) | 78 | 78 | 78 | 2 |
| Parietal endoderm | (mouse[Mesh]) AND (Embryo, Mammalian[Mesh]) AND ("Parietal endoderm"[Title/Abstract] OR "Parietal endoderms"[Title/Abstract]) | 115 | 115 | 115 | 0 |
| Pericyte | (mouse[Mesh]) AND (Embryo, Mammalian[Mesh]) AND ("Pericyte"[Title/Abstract] OR "Pericytes"[Title/Abstract]) | 58 | 58 | 58 | 13 |
| Pharyngeal | (mouse[Mesh]) AND (Embryo, Mammalian[Mesh]) AND ("Pharyngeal"[Title/Abstract] OR "Pharyngeals"[Title/Abstract]) | 119 | 119 | 119 | 24 |
| Pharyngeal epithelium | (mouse[Mesh]) AND (Embryo, Mammalian[Mesh]) AND ("Pharyngeal epithelium"[Title/Abstract] OR "Pharyngeal epitheliums"[Title/Abstract]) | 1 | 1 | 1 | 0 |
| Pineal gland | (mouse[Mesh]) AND (Embryo, Mammalian[Mesh]) AND ("Pineal gland"[Title/Abstract] OR "Pineal glands"[Title/Abstract]) | 13 | 13 | 13 | 7 |
| Platelet | (mouse[Mesh]) AND (Embryo, Mammalian[Mesh]) AND ("Platelet"[Title/Abstract] OR "Platelets"[Title/Abstract]) | 306 | 306 | 306 | 194 |
| Primordial germ cells | (mouse[Mesh]) AND (Embryo, Mammalian[Mesh]) AND ("Primordial germ cells"[Title/Abstract]) | 290 | 290 | 290 | 210 |
| Roof plate | (mouse[Mesh]) AND (Embryo, Mammalian[Mesh]) AND ("Roof plate"[Title/Abstract] OR "Roof plates"[Title/Abstract]) | 22 | 22 | 22 | 3 |
| Schwann cell | (mouse[Mesh]) AND (Embryo, Mammalian[Mesh]) AND ("Schwann cell"[Title/Abstract] OR "Schwann cells"[Title/Abstract]) | 103 | 103 | 103 | 62 |
| Sensory neuron | (mouse[Mesh]) AND (Embryo, Mammalian[Mesh]) AND ("Sensory neuron"[Title/Abstract] OR "Sensory neurons"[Title/Abstract]) | 186 | 186 | 186 | 93 |
| Spinal cord | (mouse[Mesh]) AND (Embryo, Mammalian[Mesh]) AND ("Spinal cord"[Title/Abstract] OR "Spinal cords"[Title/Abstract]) | 646 | 646 | 646 | 310 |
| Surface ectoderm | (mouse[Mesh]) AND (Embryo, Mammalian[Mesh]) AND ("Surface ectoderm"[Title/Abstract] OR "Surface ectoderms"[Title/Abstract]) | 52 | 52 | 52 | 15 |

18 Continued on next page

**Table R4 :** PubMed PMID retrieval and inclusion summary for the N4 NCBI literature datasets for the *LaManno* dataset. (Continued)

| Query label | Search query | PubMed count | Returned | Retrieved | N4 included |
|---|---|---|---|---|---|
| Undefined | (mouse[Mesh]) AND (Embryo, Mammalian[Mesh]) AND ("Undefined"[Title/Abstract] OR "Undefineds"[Title/Abstract]) | 92 | 92 | 92 | 62 |
| Vascular smooth muscle | (mouse[Mesh]) AND (Embryo, Mammalian[Mesh]) AND ("Vascular smooth muscle"[Title/Abstract] OR "Vascular smooth muscles"[Title/Abstract]) | 110 | 110 | 110 | 40 |
| Ventral hindbrain | (mouse[Mesh]) AND (Embryo, Mammalian[Mesh]) AND ("Ventral hindbrain"[Title/Abstract] OR "Ventral hindbrains"[Title/Abstract]) | 1 | 1 | 1 | 0 |
| Ventral midbrain | (mouse[Mesh]) AND (Embryo, Mammalian[Mesh]) AND ("Ventral midbrain"[Title/Abstract] OR "Ventral midbrains"[Title/Abstract]) | 30 | 30 | 30 | 0 |
| Visceral endoderm | (mouse[Mesh]) AND (Embryo, Mammalian[Mesh]) AND ("Visceral endoderm"[Title/Abstract] OR "Visceral endoderms"[Title/Abstract]) | 280 | 280 | 280 | 2 |
| Zona limitans intrathalamica | (mouse[Mesh]) AND (Embryo, Mammalian[Mesh]) AND ("Zona limitans intrathalamica"[Title/Abstract] OR "Zona limitans intrathalamicas"[Title/Abstract]) | 8 | 8 | 8 | 2 |
